# Microstructural and Functional Gradients are Increasingly Dissociated in Transmodal Cortices

**DOI:** 10.1101/488700

**Authors:** Casey Paquola, Reinder Vos De Wael, Konrad Wagstyl, Richard A.I. Bethlehem, Seok-Jun Hong, Jakob Seidlitz, Edward T. Bullmore, Alan C. Evans, Bratislav Misic, Daniel S. Margulies, Jonathan Smallwood, Boris C. Bernhardt

**Author notes:** (BCB); (CP).

## Abstract

While the role of cortical microstructure in organising neural function is well established, it remains unclear how structural constraints can give rise to more flexible elements of cognition. While non-human primate research has demonstrated a close structure-function correspondence, the relationship between microstructure and function remains poorly understood in humans, in part because of the reliance on *post mortem* analyses which cannot be directly related to functional data. To overcome this barrier, we developed a novel approach to model the similarity of microstructural profiles sampled in the direction of cortical columns. Our approach was initially formulated based on an ultra-high-resolution 3D histological reconstruction of an entire human brain and then translated to myelin-sensitive MRI data in a large cohort of healthy adults. This novel method identified a system-level gradient of microstructural differentiation traversing from primary sensory to limbic regions that followed shifts in laminar differentiation and cytoarchitectural complexity. Importantly, while microstructural and functional gradients described a similar hierarchy, they became increasingly dissociated in transmodal default mode and fronto-parietal networks. Meta analytic decoding of these topographic dissociations highlighted involvement in higher-level aspects of cognition such as cognitive control and social cognition. Our findings demonstrate a relative decoupling of macroscale functional from microstructural gradients in transmodal regions, which likely contributes to the flexible role these regions play in human cognition.

## Introduction

A core principle of neuroscience is that brain structure governs ongoing function. For decades, non-human primate work has confirmed the intrinsic relationship between microstructure and macrolevel function [1,2]. While the role of structure in cortical processing is well characterised in the sensory-motor domain, it is less clear how this constraint gives rise to more flexible elements of cognition. In an attempt to describe how cortical areas are able to take on more abstract functional roles, Mesulam (1998) postulated a hierarchical axis of large-scale cortical organisation and connectivity, referred to as the “sensory-fugal gradient” [3]. This axis describes gradual transitions at the whole-cortex level, running from primary sensory and motor regions involved in externally focussed computations towards transmodal cortices, where neural responses are not segregated by modality and operations are increasingly decoupled from perception [4,5] (see S1 Table for nomenclature). Unlike sensory cortices, transmodal cortices have a less hierarchical organisation with dense inter-connectivity and top-down projections that tend to jump synaptic levels and that allow spatially distributed areas to respond flexibly to different types of information [5–10].

The current study systematically examined the interplay between microstructural constraints and functional connectivity in humans, and its contribution to high-level cognition and behaviour. Studying cortical microstructure in humans, neuroanatomists have traditionally used cell staining techniques to map spatial variations in both cyto- and myeloarchitectural features of *post mortem* brains [11–15]. Extending upon this work, the ‘structural model’ proposes a tight coupling of microstructural similarity with increased probability of inter-areal connectivity, which was however primarily formulated on non-human primate data [1,16, see 17 for a recent review]. Although *post mortem* methods are the gold standard for describing microstructure per se, they cannot be mapped directly to function *in vivo*, making it hard to directly quantify how these metrics relate to neural function. With the advent of high-field magnetic resonance imaging (MRI), it has become possible to probe microstructural properties of different cortical regions in the living human brain. In particular, myelin sensitive imaging contrasts can differentiate regions with distinct myeloarchitectural profiles at an individual level [18–22]. In parallel, resting-state functional MRI analysis can identify highly reproducible intrinsic networks formed by cortical areas [23–28]. These studies have highlighted that transmodal cortex is largely composed of two spatially distributed yet functionally cohesive networks– the fronto-parietal network thought to respond to the demands of the moment in a flexible way [29–31] and the default mode network that depends on abstract information from self-generated memory and thought processes [32–34]. How flexible functional connectivity profiles are underpinned by microstructural differentiation remains to be seen. We expected a common sensory-fugal gradient to define microstructural and functional organisation, but we also hypothesised local differences would signify weaker hierarchical constraints in areas responsible for higher-order cognitive processes.

Core to our analysis was the formulation of a systematic approach that modelled cortico-cortical networks from similarity of microstructure profiles sampled across thousands of points and in the direction of cortical columns. The model was first developed on an ultra-high resolution *post mortem* 3D histological reconstruction of an entire human brain [35], and we show robust evidence for a principal spatial axis of gradual cytoarchitectural variation running from primary sensory to transmodal areas, recapitulating established spatial trends in laminar differentiation and cytoarchitectural complexity [3,36]. We then translated our approach to myelin-sensitive MRI in a large cohort of healthy adults, showing consistency *in vivo* and across individuals [37]. In addition to showing correspondence between histological and MRI-derived topographies, microstructure-based gradients only partially converged with macroscale functional topographies derived from task-free functional connectome analysis obtained in the same subjects [28]. In fact, while primary sensory regions served as a common anchor of microstructural and functional gradients, the microstructural axis depicted a progression towards limbic cortices, while its functional counterpart traversed towards default mode and fronto-parietal networks. Critically, meta-analytic decoding revealed that dissociation of functional from microstructural gradients was related to patterns of higher-order thought, such as a cognitive control or social cognition. Together our analyses on the convergence and divergence of spatial trends in microstructure and function support the hypothesis that reductions in hierarchical constraints in transmodal cortex is a central mechanism underlying flexible cognitive functions.

## Results

### Formulation of the histology-based microstructure profile covariance analysis

We modelled cortico-cortical microstructural similarity across a 100μm resolution Merker-stained 3D histological reconstruction of an entire *post mortem* human brain [*BigBrain*; https://bigbrain.loris.ca/main.php; [35] (Fig 1A)]. Staining intensity profiles, representing neuronal density and soma size by cortical depth, were generated along 160k surface points (*henceforth*, vertices) for each hemisphere (Fig 1B). Profile residuals, obtained after correcting intensity profile data for the *y*-coordinate to account for measurable shifts in intensity in anterior-to-posterior direction (S1 Fig), were averaged within 1012 equally sized, spatially contiguous nodes [38]. Pairwise correlations of nodal intensity profiles, covaried for average intensity profile, were thresholded at 0 and positive edges were log-transformed to produce a microstructure profile covariance matrix (MPC_HIST_); in other words, MPC_HIST_ captures cytoarchitectural similarity between cortical areas (see S2 Fig for distribution of values).

**Fig 1.**
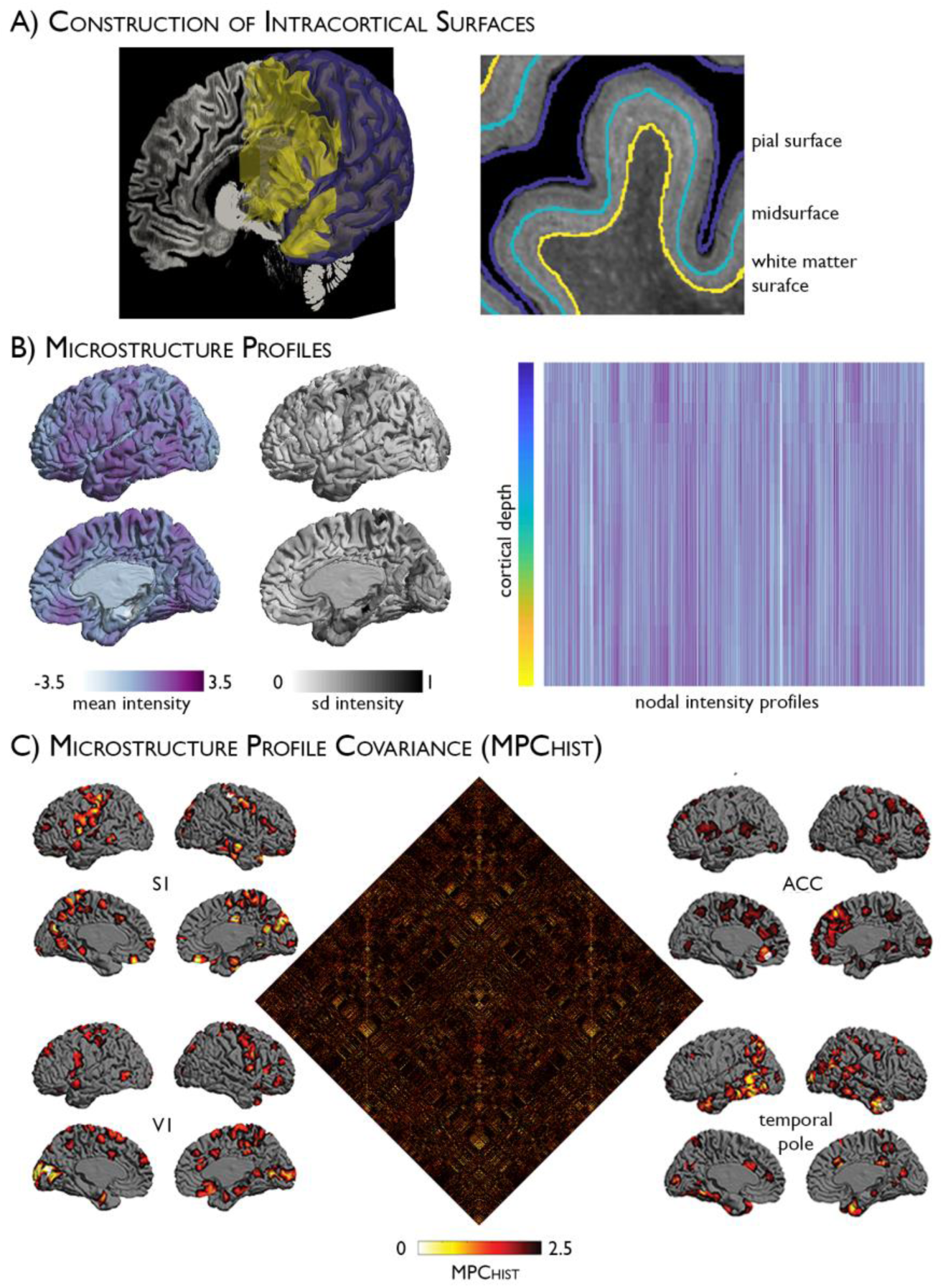
Histology-based microstructure profile covariance (MPC_HIST_) analysis. **(A)** Pial *(purple)* and white matter *(WM: yellow)* surfaces displayed against a sagittal slice of the BigBrain *(left)* and with the midsurface *(blue)* in magnified view *(right)*. (**B**) Mean and standard deviation (SD) in residual intensity at each node are displayed on the cortex *(left)*. Cortex-wide intensity profiles were calculated by systematic intensity sampling across intracortical surfaces (*rows*) and nodes (*columns*). (**C**) The MPC_HIST_ matrix depicts node-wise partial correlations in intensity profiles, controlling for the average intensity profile. Exemplary patterns of microstructural similarity from primary somatosensory (S1), anterior cingulate cortex (ACC), primary visual (V1) and the temporal pole. Seed nodes are shown in *white*. Histological data is openly available as part of the Big Brain initiative (https://bigbrain.loris.ca/main.php).

The pipeline was optimised with respect to the number of intracortical surfaces based on matrix stability (see Methods and S3 Fig). While microstructural similarity had a small but significant statistical relationship with spatial proximity (adjusted R^2^=0.02, p<0.001), similar findings were obtained after correcting for geodesic distance.

### The principal gradient of microstructural similarity reflects sensory-fugal neurostructural variation

Diffusion map embedding, a nonlinear dimensionality reduction algorithm [39] recently applied to identify an intrinsic functional segregation of cortical regions based on resting state functional MRI [28], was applied to the histology-based microstructure profile covariance matrix (Fig 2A). The relative positioning of nodes in this embedding space informs on (dis)similarity of their covariance patterns. The first principal gradient (G1_HIST_), accounting for 14.5% of variance, was anchored on one end by primary sensory and motor areas and on the other end by transmodal association and limbic cortices (Fig 2B; see S4 Fig for the second gradient and S5 Fig for results on inflated cortical surfaces). G1_HIST_ depicted the most distinguishable transition in the shape of microstructure profiles (Fig 2B Right). Regions of the prefrontal cortex *(green)* expressed an intracortical profile that was closest to the cortex-wide average. Extending outwards from centre of G1_HIST_, sensory and motor regions *(blue-purple)* exhibited heightened cellular density around the midsurface, whereas paralimbic cortex *(red)* displayed specifically enhanced density near the cortical borders. For further validation of the biological basis of G1_HIST_, we mapped independent atlases of laminar differentiation [40] and cytoarchitectural class [13,41] onto the BigBrain midsurface (Fig 2C).

**Fig 2.**
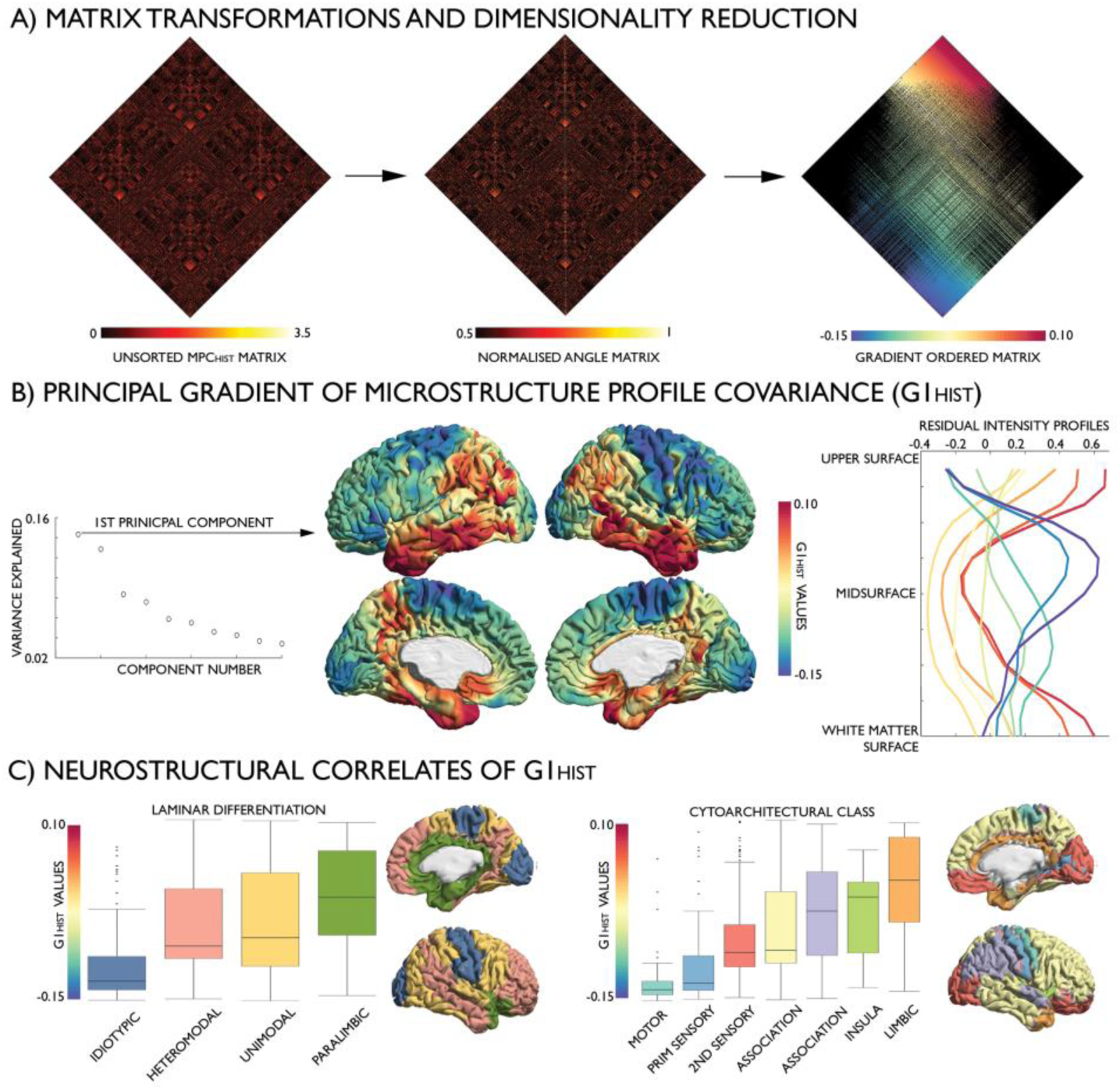
The principal gradient (G1_HIST_) of the histology-based microstructure profile covariance matrix (MPC_HIST_). **(A)** Identification: the MPC_HIST_ matrix was transformed into an affinity matrix, which captures similarities in microstructural profile covariance between nodes; this affinity matrix was subjected to diffusion map embedding, a non-linear compression algorithm that sorts nodes based on MPC_HIST_ similarity. **(B)** Variance explained by embedding components (*left*). The first component, G1_HIST_, describes a gradual transition from primary sensory and motor (*blue*) to transmodal and limbic areas (*red*), corresponding to changes in intensity profiles, illustrated with the mean residual intensity profile across ten discrete bins of the gradient (*right)*. **(C)** Spatial associations between G1_HIST_ and levels of laminar differentiation [*left;* [40]] and cytoarchitectural taxonomy [*right;* [13,41]], ordered by median. Histological data is openly available as part of the Big Brain initiative (https://bigbrain.loris.ca/main.php).

Multiple linear regression analyses showed that levels of laminar differentiation and cytoarchitectural taxonomy each accounted for 17% of variance in G1_HIST_ (S2-3 Tables). Strongest predictors were idiotypic (β=0.06, p<0.001) and paralimbic (β=0.05, p<0.001) laminar differentiation levels, and limbic (β=0.07, p<0.001) as well as motor (β=0.06, p<0.001) classes in the cytoarchitectural model, demonstrating the cytoarchitectural distinctiveness of regions at the extremes of G1_HIST_.

### Development of in vivo microstructure profile covariance analysis

The microstructure profile covariance approach was adapted to *in vivo* data in single individuals using T1w/T2w MRI, a ratio indexing cortical microstructure and myelin [18], shown to recapitulate sensory-fugal transitions [42]. Multimodal surface-matched T1w/T2w images, pial surfaces, and white matter surfaces were obtained from the minimally-preprocessed S900 release of the Human Connectome Project [37,43]. We selected a total of 219 unrelated subjects and grouped these randomly into a *Discovery* sample (n=110, 66 females, mean±SD age=28.8±3.8 years) and a *Replication* sample (n=109, 62 females, mean±SD age=28.5±3.7 years). For each individual, we systematically generated intracortical surfaces using the same equivolumetric transformation algorithm as for the histological dataset [44,45], and aggregated whole-cortex intensity profiles across 64,984 linked vertices that were subsequently parcellated into 1012 contiguous nodes [38]. We computed pairwise partial correlations between nodal intensity profiles (controlling for the average intensity profile), kept only positive correlations, and log-transformed the result to produce a cortex-wide microstructure profile covariance matrix (MPC_MRI_: see S6 Fig for distribution of values). A group-average MPC_MRI_ matrix was calculated across all participants in the *Discovery* sample. While microstructural similarity estimated *in vivo* was stronger between proximal nodes, the variance accounted for by geodesic distance was low (adjusted R^2^ = 0.04, β=-2.57, p<0.001).

Diffusion map embedding revealed a principal gradient of microstructural differentiation accounting for 13.7% of variance (G1_MRI_, Fig 3B; see S7 Fig for the second gradient). In line with its histological counterpart, G1_MRI_ was anchored on one end by primary sensory areas and on the other end by limbic regions. Cortex-wide analysis demonstrated a high correlation of G1_HIST_ and G1_MRI_ at a group level (r=0.63, p<0.001) as well as an individual level (0.41<r<0.62, mean r=0.52±0.03), driven by the close spatial correspondence of gradient extremes (Fig 3C, *left*), and was unrelated to inter-individual variance in cortical curvature (r=-0.01, p=0.867). G1_MRI_ depicts increasing mean myelin content, as well as a gradual transition in the relative myelin content around the midsurface (Fig 3B, *right*). Microstructural profiles of the prefrontal cortex *(orange)* again resembled the cortex-wide average; however, comparing node ranks across both modalities revealed a shift in prefrontal regions towards the transmodal anchor in G1_MRI_ (Fig 3C, *right*). This effect appeared to be driven by a downward shift of lateral occipital-parietal areas towards the sensory anchor, owing to heavy myelination relative to their cytoarchitectural complexity [46]. Laminar differentiation and cytoarchitectural taxonomy accounted for 44% and 37% of variance in G1_MRI,_ respectively (S4-5 Tables). As in the histological analysis, the paralimbic (β=0.10, p<0.001) and idiotypic (β=0.07, p<0.001) levels were the strongest predictors within the laminar differentiation model, while motor (β=0.15, p<0.001), limbic (β=0.07, p<0.001), and primary sensory (β=0.10, p<0.001) classes were strong predictors within the cytoarchitectural model.

**Fig 3.**
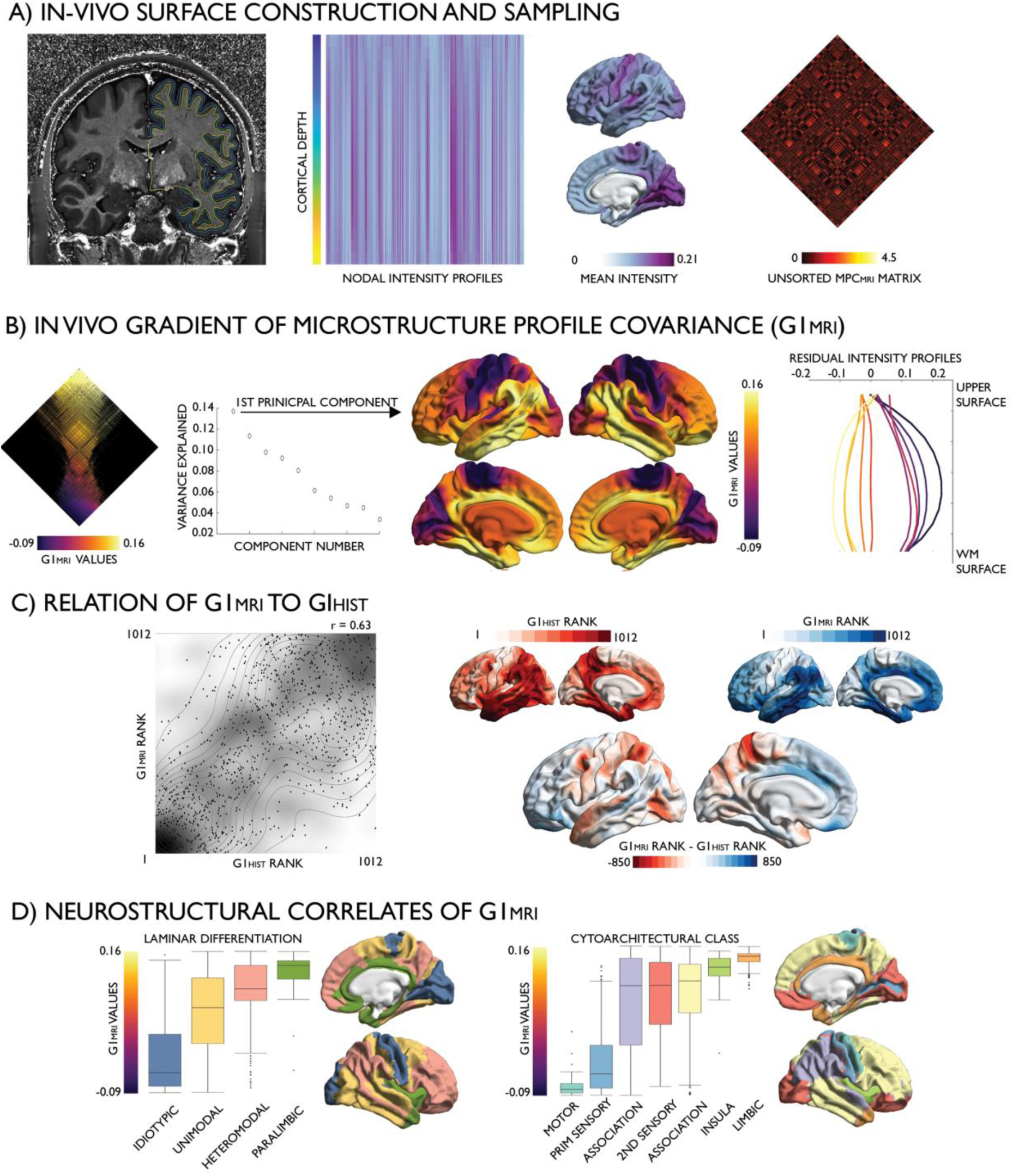
*In vivo* microstructure profile covariance (MPC_MRI_). **(A)** Left hemisphere pial, mid, and white matter surfaces superimposed on a T1w/T2w image (*left*); whole-cortex intensity profiles were calculated by systematic sampling across surfaces (*rows*) and vertices, and then averaged with each node (*columns*). Mean at each node (*centre right*); MPC_MRI_ matrix depicts node-wise partial correlations in intensity profiles, covaried for mean whole-cortex intensity profile *(right)*. **(B)** Normalised angle matrix sorted by the principal gradient (*left*); variance explained by the diffusion embedding components (*left centre*) and the principal gradient (*right centre*); mean residual intensity profiles within ten discrete bins of the gradient (*right).* **(C)** Similarity of histological and *in vivo* gradients (G1_HIST_, G1_MRI_) shown in a density plot (*left)* and node-wise rank differences shown on the cortical surfaces (*right).* **(D)** Associations of G1_MRI_ to levels of laminar differentiation [40] and cytoarchitectural class [13,41] ordered by median. In-vivo imaging data is openly available as part of the Human Connectome Project S-900 release (https://www.humanconnectome.org/study/hcp-young-adult/document/900-subjects-data-release).

### Correspondence of microstructural similarity with macroscale functional organisation

To examine the role of microstructural similarity in macroscale functional organisation, we generated a group-average resting state functional connectome across the *Discovery* subsample and derived gradients with diffusion map embedding. As shown previously [28], the principal functional gradient (G1_FUNC_) extends from primary sensory and motor networks, through dorsal attention and salience networks, to finally culminate in the transmodal core composed of fronto-parietal and default mode networks (Fig 4A). At the group level, cortex-wide analyses demonstrated moderate-to-high correlations of G1_MRI_ with G1_FUNC_ (r=0.52, p<0.001; Fig 4B) and G1_HIST_ with G1_FUNC_ (r=0.31, p<0.001), illustrating a common topography of microstructural profile covariance and functional connectivity. Additionally, the topographies of microstructure and function were more closely related than node-to-node correspondence of microstructural similarity with functional connectivity (*in vivo:* r=0.10, p<0.001; histology-based: r=0.11, p<0.001), supporting the utility of connectivity-informed dimensionality reduction techniques to reveal common principles of sensory-fugal cortical organisation [47].

**Fig 4.**
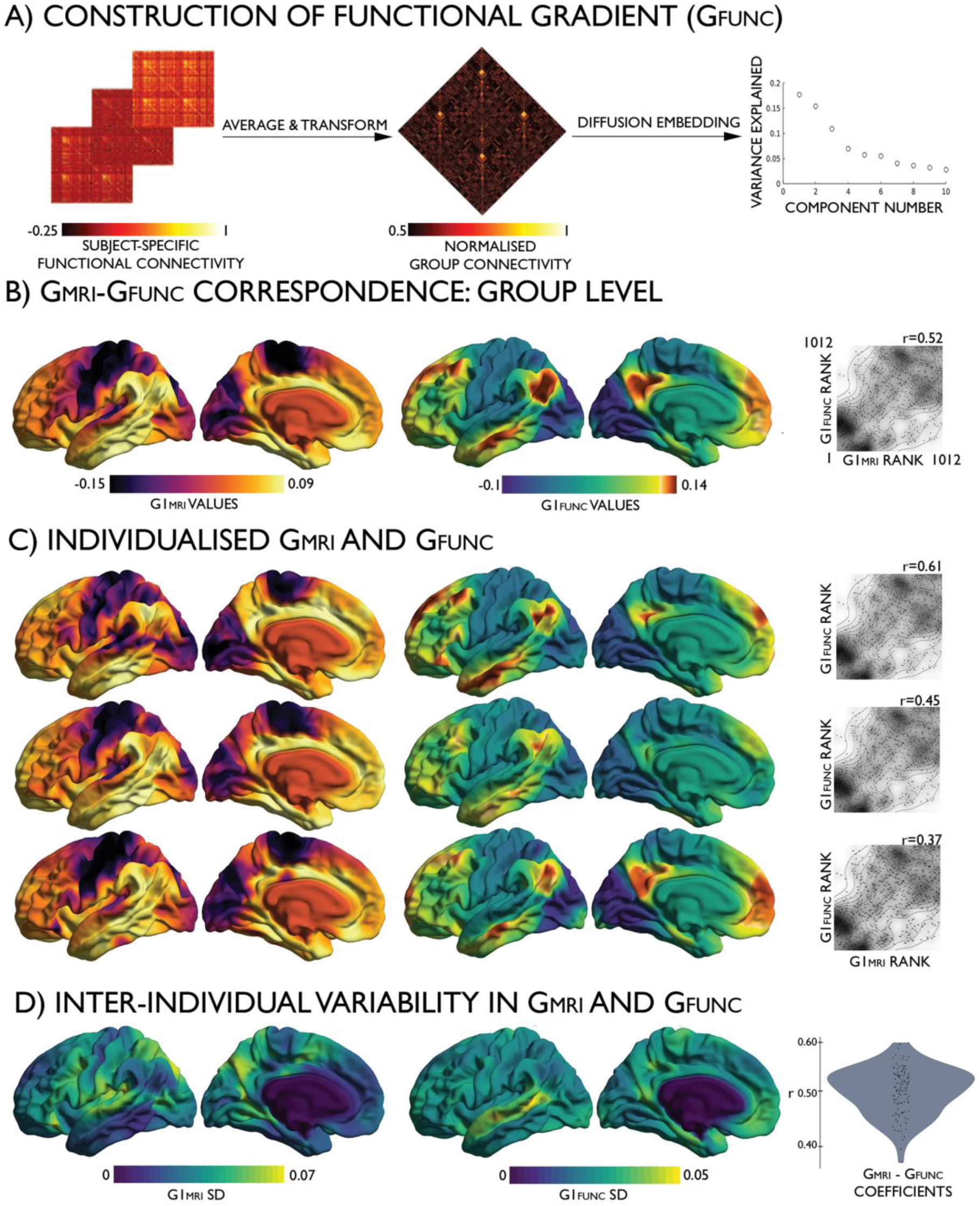
Cross-modal correspondence of the MPC_MRI_ and intrinsic functional gradients. (A) Transformation from individual functional connectomes (*left)* to a group average normalised angle matrix (*centre)* to diffusion embedding components *(right)* (B) The group level G1_MRI_ *(left)*, group level G1_FUNC_ (*centre)* and density plots depicting the correlation between the gradients (*right)*. (C) Consistency across 3 example subjects and (D) inter-individual variability of the gradients and cross-modal correspondence in the *Replication* dataset. In-vivo imaging data is openly available as part of the Human Connectome Project S-900 release (https://www.humanconnectome.org/study/hcp-young-adult/document/900-subjects-data-release).

The *Replication* dataset underwent identical processing procedures as the *Discovery* dataset. At a group level, G1_MRI_ derived from the *Replication* and *Discovery* cohorts were highly correlated (r=0.98, p<0.001). High correlations between G1_MRI_ and G1_FUNC_ were also evident at the individual subject level, following alignment of individual functional and microstructural gradients to templates built from the group *Discovery* dataset (see Methods, for details). For every participant, we observed a significant correlation between G1_MRI_ and G1_FUNC_ (0.37<r<0.61, p<0.001; Fig 4C and 4D).

In addition to studying their commonalities, we assessed unique topographic features of the modality-specific gradients. Cortex-wide nodal rank comparisons highlighted an upward shift in the position of the prefrontal cortex and precuneus in functional gradient space, relative to the microstructure-derived gradient, and a converse downward shift of the posterior inferior temporal and midcingulate cortices (Fig 5A, see S8 Fig for replication at vertex-level). These shifts were independent of regional differences in the inter-individual variance of curvature (r=0.01, p=0.741). Notably, the greatest rank differences were evident in transmodal cortices, suggesting a specific dissociation of function from microstructure in these higher-order regions. Inspecting the distribution of nodes in intrinsic functional communities [26] along each gradient (Fig 5B), we noted that while the sensory-fugal gradient was overall preserved across all modalities, different sensory and transmodal networks occupied extremes in each modality. G1_MRI_ extended from somatomotor to limbic networks, whereas G1_FUNC_ extended from visual to transmodal default mode networks. Comparing the average node rank of each functional community, between modalities and across individuals (S6 Table), we noted that the default mode and fronto-parietal networks shifted to the apex of G1_FUNC,_ reflecting segregation of higher-order communities during rest, despite their similar myeloarchitecture.

**Fig 5.**
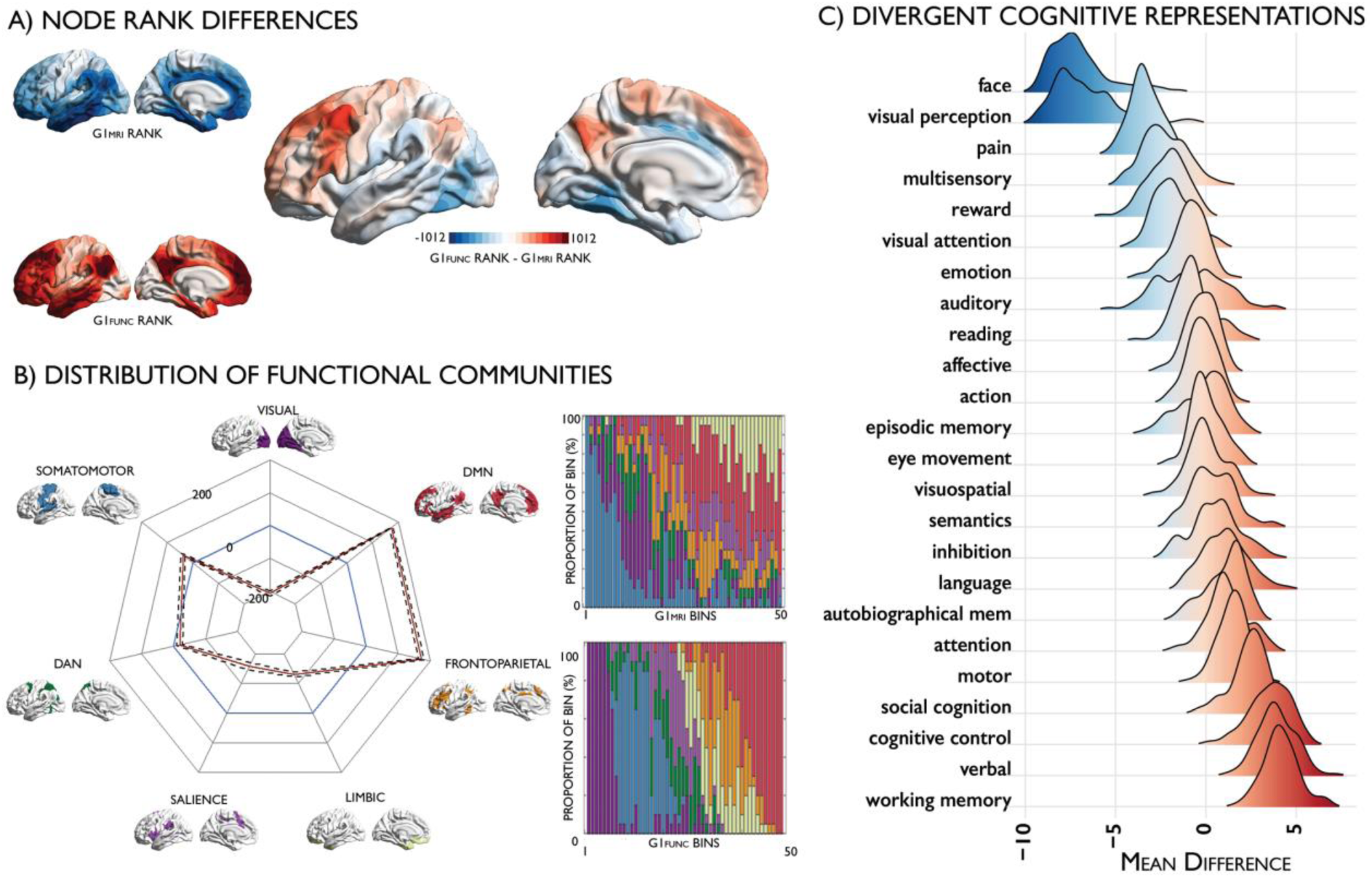
Divergent representations of the cortical hierarchy derived from microstructure and function. (A) Differences in nodal ranks between G1_MRI_ *(blue)* and G1_FUNC_ *(red).* (B) Radar plot depicting the difference in mean node ranks of functional communities [26] between G1_MRI_ *(blue)* and G1_FUNC_ *(red)*, with 95% confidence intervals calculated across individuals presented with dotted lines; stacked bar plots depicting the proportion of each bin accounted for by intrinsic functional communities. (C) Meta-analysis maps for diverse cognitive terms were obtained from Neurosynth [28]. We calculated node-wise z-statistics, capturing node-term associations, and calculated the centre of gravity of each term along G1_FUNC_ and G1_MRI._ The density plots depict the mean difference in the centre of gravity of meta-analysis maps in G1_FUNC_ and G1_MRI_ space across subjects. In-vivo imaging data is openly available as part of the Human Connectome Project S-900 release (https://www.humanconnectome.org/study/hcp-young-adult/document/900-subjects-data-release).

Our final analysis examined whether the functional topography diverged from microstructure specifically in cortical areas involved in abstract, perceptually decoupled functions. We conducted a meta-analysis using the Neurosynth database and estimated the centre of gravity across a set of diverse cognitive terms [28] along G1_FUNC_ relative to G1_MRI_ (S5c Fig). Top terms exhibiting the strongest upward shift from G1_MRI_ to G1_FUNC_ involved multi-domain, integrative functions, such as “working memory”, “verbal”, “cognitive control” and “social control”, whereas the strongest downward shifts involved higher-order visual processing, such as “face” and “visual perception”.

### Robustness of the MPC and resultant gradients

#### A series of replication analyses assessed the robustness of the MPC framework

##### a) Histology

G1_HIST_ was highly consistent across variations in thresholding, parcellation, intracortical surface number, and BigBrain voxel resolution (S3 Fig). Furthermore, varying the α parameter for diffusion map embedding algorithm from 0 and 1 (in increments of 0.1) resulted in virtually identical gradients (all r>0.99). We also compared diffusion map derived gradients to more conventional graph theoretical characterisations of the MPC_HIST_ matrix. Specifically, we applied Louvain modularity detection [48], incorporating a procedure that maximized consensus across fine- and coarse-grained decompositions (by varying the tuning parameter γ from 0.5-1.5) across 100 repetitions. We resolved three modules (0.49<Q<0.71; S9 Fig), which occupied distinct positions along the first two gradients. Diffusion map embedding thus represents a continuous and complementary approach to order microstructure profile covariance, which emphasises gradual transitions within and between discrete cortical areas that are fundamental to the hierarchical organisation of layer-specific cortical projections [49].

##### b) In vivo

As for their histological counterpart, G1_MRI_ solutions were robust against variations in processing parameters, including matrix thresholding, parcellation scheme, and surface number (S10 Fig). Again, variation of the α parameter resulted in virtually identical gradients (all r>0.99). Application of Louvain community detection identified only two modules in the microstructure profile covariance matrix which concisely halved G1_MRI_ (0.35<Q<0.41, S11 Fig). We replicated G1_MRI_ in an independent dataset of healthy late adolescents/young adults [50] that underwent magnetisation transfer imaging (spatial correlation: r=0.78, p<0.001, S12 Fig) and in a cohort of 17 healthy adults scanned at our imaging centre in whom quantitative T1 relaxation data was available (spatial correlation: r=0.81, p<0.001, S13 Fig), demonstrating robustness of the approach to acquisition site, acquisition type, and surface construction.

## Discussion

Cortical areas identified by classical neuroanatomical studies represent discrete regions embedded within gradual transitions in cyto- and myeloarchitecture [11–13,51]. Cortical gradients are nearly ubiquitous across microstructural and functional domains of the mammalian neocortex, where mounting evidence supports a common and overarching “sensory-fugal” organisation [49,52]. Capitalizing on a ultra-high-resolution 3D reconstruction of an entire *post mortem* human brain [35], we sampled microstructure profiles and utilised unsupervised techniques to identify smooth transitions in cytoarchitectural composition. We translated our approach from histology to myelin-sensitive *in vivo* MRI and recovered similar microstructural gradients, which were consistent across individuals. Collectively, our findings support a common sensory-fugal axis of inter-areal cortical differentiation across histology and *in vivo* microstructure. Importantly, we also observed a divergence between microstructural and functional gradients towards transmodal default mode and fronto-parietal networks. While these functional networks were distributed across the microstructural gradient, they were localised to the top of the functional gradient. Based on meta-analytical decoding, this divergence was found to relate to aspects of human cognition such as cognitive control and social cognition, supporting views that these regions may be relatively untethered from hierarchical constraints that allow them to take on more flexible cognitive roles.

To bridge different scales of human brain organisation [53], we developed a cortico-cortical network model based on microstructural similarity between areas. Microstructure profile generation, based on state-of-the-art equivolumetric surface construction techniques [44], provided a simplified recapitulation of cellular changes across the putative laminar structure of the cortex. Importantly, our covariance framework generated cytoarchitecturally grounded networks, which were sensitive to laminar thickness as well as cell size and density. The current study revealed gradual cytoarchitectural transitions across the folded neocortex with primary sensory areas on one end of the gradient, and cingulate and parahippocampal cortices on the other end. These architectonically unique regions anchor the observed macroscopic sensory-fugal axis of microstructural differentiation, reflecting a continuum through distinct levels of laminar differentiation and cytoarchitectural complexity, from the koniocortical primary sensory areas, via unimodal association areas towards dysgranular paralimbic cortices [40,54]. The findings strongly support prior evidence on the successive steps of change in cortical architecture and connectivity in reptiles [55], monotremes [56], marsupials [57,58], cetaceans [59], prosimians [60], squirrel monkeys [61], rhesus monkeys [4,62,63], and humans [36,49,64]. Despite differences in cytoarchitectural nomenclature across prior research (see also S1 Table), our findings also mirror the watercolour illustrations of cortical gradients created by Bailey and von Bonin (1951), which highlight the distinctiveness of these gradient anchors.

We directly translated our approach to myelin sensitive *in vivo* MRI [18–22], recovering highly consistent topographic maps. Despite there not being a universal 1:1 mapping between cyto- and myeloarchitecture, they are closely related and complimentary aspects of the same cortical wiring system [14,65,66]. The size, density and position of pyramidial neurons determine the total intracortical myelin content, as well as myelin banding. Histological studies have shown how this micro-scale correspondence confer similar macro-scale topographies, whereby laminar differentiation and mean intracortical myelin gradually change along the sensory-fugal gradient [54,67–69]. Our findings extend upon this work by showing the global topography of myeloarchitectural similarity, which involves both mean myelin and myelin banding [66], is strongly related to macro-scale cytoarchitectural differentiation.

Microstructural and functional gradients were consistently anchored by primary sensory regions on the lower end, recapitulating their consistent location at the bottom of functional and structural hierarchies in primates [3,7,28,36,70,71]. In primary regions, where neural processes are strongly constrained by sensorimotor contingencies, information processing is driven by extrinsic inputs and interactions with the outside world [72,73], as well as intrinsic signalling molecules involved in axonal guidance, cell adhesion, regional circuit formation, and thus cortical patterning [74]. The impact of these forces decreases with synaptic distance, producing a sensory-fugal axis of neurostructural and functional differentiation along the cortical mantle. In fact, given the default mode network’s location as the most distant functional system relative to the location of primary sensory sulci [28], it may be relatively untethered by the influence of intrinsic signalling molecules and extrinsic activity [75], and thus expresses less unique microstructural profiles, as observed here and in classical neuroanatomical studies [11–13].

By showing a close correlation between individualised microstructural gradients and corresponding functional connectome gradients [28], our findings expand the structural model of connectivity to living humans. Originally formulated in non-human primates [1,2] and later demonstrated in rodents [76] and cats [77,78], the structural model predicts increased projections between areas with more similar internal architecture. This principle underpins concomitant variations in cytoarchitecture and patterns of connectivity along the sensory-fugal axis and, despite providing a parsimonious account of cortical connectivity, had not been previously explored in living humans or using functional connections. A recent study carried out diffusion MRI network construction and showed an association between structural connectivity strength in selected regions to microstructural similarity information obtained from BigBrain [79]. While this study demonstrated the feasibility of integrating BigBrain data with *in vivo* connectomics, our study had a different scope. Firstly, we formulated a novel step-by-step procedure to derive networks of microstructural similarity, both based on BigBrain as well as microstructural MRI, bringing a new network modality to connectomics. Of note, MPC networks have whole-cortex coverage and their construction accounts for curvature using state-of-the-art equivolumetric surface transformation techniques. As we have shown, the MPC procedure can be consistently translated from histology to high resolution *in vivo* imaging, thus allowing a direct comparison of microstructural similarity networks with those based on functional connectivity, and thus probes structure-function associations in the same subjects.

In contrast to a close functional coupling of microstructurally similar areas at the low end of the cortical gradient, the functional connectivity between higher-order regions may not be as strongly constrained by hierarchical principles. For example, prefrontal and parietal association regions bilaterally project feedforward and feedback connections [80], which can be densely interdigitated [81]. The diverse connectivity profiles in transmodal cortices are likely related to heightened synaptic plasticity [69] that enables more flexible reconfigurations of functional relationships. In contrast, synaptic plasticity is lower in sensory cortices where quick, accurate response to external stimuli necessitate a constrained hierarchical organisation [3]. Another plausible mechanism for reduced influence of microstructural homophily in higher-order networks may be the increased relevance of long-range structural connections [82]. Such wider ranging structural connections may accommodate the more distributed spatial layout and diverse functional roles of transmodal cortices. Conversely, primary sensory and motor regions exhibit more locally clustered short range connectivity profiles related to microstructural similarity [83,84], likely in accordance with their more specialized and stable functional roles.

Our approach was robust with respect to variations in algorithmic and analytical choices. While the singular nature of the BigBrain dataset prohibited replication of the histological pipeline, we demonstrated consistency of neuroimaging-derived microstructural gradients across three cohorts with unique myelin sensitive contrasts [18,20,22,37,50]. Importantly, the *in vivo* approach can serve as a lower resolution, yet biologically meaningful extension of the histological work. In fact, myelin-based gradients exhibited a comparable association with laminar differentiation and cytoarchitectural complexity as the histology-based gradient, as shown to be the case by earlier *post mortem* work [66–69]. Albeit replication of our findings at a vertex-wise level, the millimetre resolution of *in vivo* imaging may not always capture detailed spatial features visible on histology. With the emergence of algorithms that detect cortical laminae based on histological data [85] and increasing availability of ultra-high field MRI scanners (at field strengths of 7T and higher), we expect that the histological and *in vivo* MPC approach may be further refined in future work, providing an even more direct bridge between the micro- and macro-scales of human brain organisation. As it offers a hierarchy-dependent reference frame to examine the interplay of microstructural and system-level network mechanisms in single individuals, the proposed framework may also be advantageous in neurodevelopmental research and complement existing covariance network mapping approaches that have shown promise in studying typical and atypical development [86–91]. As such, it offers a powerful and freely available (http://github.com/micaopen/MPC) tool to investigate coordinated changes in cortical microstructure paralleling the emergence and maturation of large-scale functional networks during development [92,93], and may conversely provide insight into the structural underpinnings of atypical network configurations in complex neurodevelopmental disorders [94].

We close by considering the significance of the observed progressive dissociation between structural and functional topographies in cortical areas involved in multi-domain, integrative processing. Components of the fronto-parietal network have been shown to guide behaviour in an adaptive manner, changing in line with external demands [29], and possibly substantiated by its ability to dynamically reconfigure its functional connectivity [95]. Although the functional role of the default mode network as the putative apex of the cortical functional hierarchy in primates remains subject to debate [96], it is evidently involved a broad class of memory-driven operations, involving self-referential and simulative thought processes and some degree of abstraction [32–34]. The interesting possibility that reduced hierarchical constraints enable functional diversity and flexibility was also supported by *ad hoc* meta-analysis, suggesting that microstructural and functional gradients decouple in regions contributing to processes such as working memory, social cognition, and cognitive control. Such a hypothesis provides a potential mechanistic account for why some of the more creative acts of the human mind emerge through the interaction of the two most dominant yet structurally diverse functional systems [97].

## Acknowledgements

The authors would also like to express their gratitude to the teams at the Foschungszentrum Julich and the Montreal Neurological Institute who made the BigBrain dataset available. Furthermore, we thank Dr Nicola Palamero-Gallagher for helpful and inspiring discussions.

## Methods

### Histology-based MPC

#### Histological data acquisition and preprocessing

An ultra-high resolution Merker stained 3D volumetric histological reconstruction of a *post mortem* human brain from a 65-year-old male was obtained from the open-access BigBrain repository on February 2, 2018 [https://bigbrain.loris.ca/main.php; [35]]. The *post mortem* brain was paraffin-embedded, coronally sliced into 7,400 20μm sections, silver-stained for cell bodies [98] and digitised. Manual inspection for artefacts (*i.e.*, rips, tears, shears, and stain crystallisation) was followed by automatic repair procedures, involving non-linear alignment to a *post mortem* MRI, intensity normalisation, and block averaging [99]. 3D reconstruction was implemented with a successive coarse-to-fine hierarchical procedure [100]. We downloaded the 3D volume at four resolutions, with 100, 200, 300, and 400μm isovoxel size. We primarily analysed 100μm data and used 200, 300 and 400μm data to assess consistency of findings across spatial scales. Computations were performed on inverted images, where staining intensity reflects cellular density and soma size. Geometric meshes approximating the outer and inner cortical interface (*i.e.*, the GM/CSF boundary and the GM/WM boundary) with 163,842 matched vertices per hemisphere were also available [101].

#### Histology-based microstructure profile covariance (MPC) analysis

##### a) Surface sampling

We systematically constructed 10-100 equivolumetric surfaces, in steps of 1, between the outer and inner cortical surfaces [45]. The equivolumetric model compensates for cortical folding by varying the Euclidean distance, ρ, between pairs of intracortical surfaces throughout the cortex to preserve the fractional volume between surfaces [44]. ρ was calculated as follows for each surface

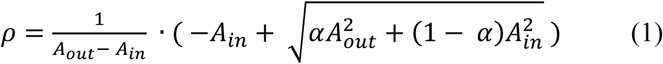

where α represents fraction of the total volume of the segment accounted for by the surface, while A_out_ and A_in_ represent the surface area of the outer and inner cortical surfaces, respectively. Next, vertex-wise microstructure profiles were estimated by sampling intensities along linked vertices from the outer to the inner surface across the whole cortex. In line with previous work [85], layer 1 was approximated as the top ten percent of surfaces and removed from the analysis due to little inter-regional variability. Note however, that findings were nevertheless virtually identical when keeping the top ten percent of surfaces. To reduce the impact of partial volume effects, the deepest surface was also removed. Surface-based linear models, implemented via SurfStat for Matlab (http://mica-mni.github.io/surfstat) [102], were used to account for an anterior-posterior increase in intensity values across the BigBrain due to coronal slicing and reconstruction [35], whereby standardised residuals from a simple linear model of surface-wide intensity values predicted by the midsurface y-coordinate were used in further analyses.

##### b) MPC matrix construction

Cortical vertices were parcellated into 1,012 spatially contiguous cortical ‘nodes’ of approximately 1.5cm^2^ surface area, excluding outlier vertices with median intensities more than three scaled median absolute deviations away from the node median intensity. The parcellation scheme preserves the boundaries of the Desikan Killany atlas [38] and was transformed from conte69 surface to the BigBrain midsurface via nearest neighbour interpolation. Nodal intensity profiles underwent pairwise Pearson product-moment correlations, controlling for the average whole-cortex intensity profile. MPC_HIST_ for a given pair of nodes *i* and *j* was thus

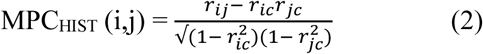

where *r*_*ij*_ is the Pearson product moment correlation coefficient of the BigBrain intensity profiles at nodes *i* and *j, r*_*ic*_ the correlation coefficient of the intensity profile at node *i* with the average intensity profile across the entire cortex, and *r*_*jc*_ the Pearson correlation of the intensity profile at node *j* with the average intensity profile across the whole brain. The MPC matrix was thresholded above zero and remaining MPC values were log-transformed to produce a symmetric 1012×1012 MPC_HIST_ matrix. The in-house developed code for MPC construction is available online (https://github.com/MICA-MNI/micaopen/MPC).

##### c) Parameter estimation

The optimal surface number was determined based on the stability of the MPC matrix. This procedure involved (repeatedly and randomly) dividing the vertex intensity profiles within each node into two groups and constructing two MPC matrices, then calculating the Euclidean distance between them. The procedure was repeated 1000 times. Although the MPC matrix instability was robust to variations in surface number, the 18-surface solution exhibited a noticeable local minimum MPC instability in the studied range (10-100 surfaces) and was used in subsequent analyses (**S2 Fig**). Notably, the MPC gradient was similar using two finer grained solutions (*i.e.*, 54 and 91 surfaces), where local minima were observed as well. More details on the origins of the stability statistic in clustering algorithms may be found elsewhere [103].

##### d) Relation to spatial proximity

To determine whether MPC_HIST_ was not purely driven by spatial proximity, we correlated MPC_HIST_ strength with geodesic distance for all node pairs. The latter was calculated using the Fast Marching Toolbox between all pairs of vertices, then averaged by node (https://github.com/gpeyre/matlab-toolboxes/tree/master/).

#### Histology-based MPC gradient mapping

In line with previous studies [28,104], the MPC_HIST_ matrix was proportionally thresholded at 90% per row, and converted into a normalised angle matrix. Diffusion map embedding [39], a nonlinear manifold learning technique, identified principal gradient components explaining MPC_HIST_ variance in descending order (each of 1×1012). In brief, the algorithm estimates a low-dimensional embedding from a high-dimensional affinity matrix. In this space, cortical nodes that are strongly inter-connected by either many suprathreshold edges or few very strong edges are closer together, whereas nodes with little or no inter-covariance are farther apart. The name of this approach, which belongs to the family of graph Laplacians, derives from the equivalence of the Euclidean distance between points in the embedded space and the diffusion distance between probability distributions centred at those points. Compared to other non-linear manifold learning techniques, the algorithm is relatively robust to noise and computationally inexpensive [105,106]. Notably, it is controlled by a single parameter α, which controls the influence of the density of sampling points on the manifold (α = 0, maximal influence; α = 1, no influence). In this and previous studies [28,104], we followed recommendations and set α=0.5, a choice that retains the global relations between data points in the embedded space and has been suggested to be relatively robust to noise in the covariance matrix. Gradients were mapped onto BigBrain midsurface visualised using SurfStat (http://mica-mni.github.io/surfstat) [102], and we assessed the amount of MPC_HIST_ variance explained. To show how the principal gradient in MPC_HIST_ (G1_HIST_) relates to systematic variations in microstructure, we calculated and plotted the mean microstructure profiles within ten equally-sized discrete bins of G1_HIST_.

#### Relation of G1_HIST_ to laminar differentiation and cytoarchitectural taxonomy

We evaluated correspondence of G1_HIST_ to atlas information on laminar differentiation and cytoarchitectural class. To this end, each cortical node was assigned to one of four levels of laminar differentiation (*i.e.*, idiotypic, unimodal, heteromodal, paralimbic) derived from a seminal model of Mesulam, which was built on the integration of neuroanatomical, electrophysiological, and behavioural studies in human and non-human primates [40], and one of the seven Von-Economo/Koskinas cytoarchitectural classes (*i.e.*, primary sensory, secondary sensory, motor, association 1, association 2, limbic or insular) [13,41]. In the case of laminar differentiation maps, assignment was done manually; in the case of cytoarchitectural classes, we mapped previously published Von Economo/Koskinas classes [91] to the BigBrain midsurface with nearest neighbour interpolation and assigned nodes to the cytoarchitectural class most often represented by the underlying vertices. Finally, we estimated the contribution of level of laminar differentiation (D, a categorical variable) and cytoarchitectural class (C, a categorical variable) to the principal gradient G1_HIST_ of the MPC_HIST_ within two separate multiple regression models:

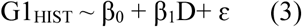

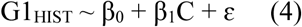

We evaluated model fit via adjusted R^2^ statistics and unique variances explained by each predictor (β).

### In vivo MPC

#### MRI data acquisition and preprocessing

We studied data from 219 unrelated healthy adults from the minimally preprocessed S900 release of the Human Connectome Project (HCP; Glasser et al., 2013). The *Discovery* dataset included 110 individuals (66 females, mean±SD age=28.8±3.8 years) and the *Replication* dataset 109 (62 females, mean±SD age=28.5±3.7 years). MRI data were acquired on the HCP’s custom 3T Siemens Skyra equipped with a 32-channel head coil. Two T1w images with identical parameters were acquired using a 3D-MPRAGE sequence (0.7mm isotropic voxels, matrix=320×320, 256 sagittal slices; TR=2400ms, TE=2.14ms, TI=1000ms, flip angle=8°; iPAT=2). Two T2w images were acquired using a 3D T2-SPACE sequence with identical geometry (TR=3200ms, TE=565ms, variable flip angle; iPAT=2). Four rs-fMRI scans were acquired using multi-band accelerated 2D-BOLD echo-planar imaging (2mm isotropic voxels, matrix=104×90, 72 sagittal slices; TR=720ms, TE=33ms, flip angle=52°; mb factor=8; 1200 volumes/scan). Participants were instructed to keep their eyes open, look at fixation cross, and not fall asleep. While T1w and T2w scans were acquired on the same day, rs-fMRI scans were split over two days (two scans/day).

Structural and resting state functional MRI data underwent HCP’s minimal preprocessing [43,107,108]. For structural MRI, images underwent gradient nonlinearity correction. When repeated scans were available, these were co-registered and averaged. Following brain extraction and readout distortion correction, T1w and T2w images were co-registered using rigid body transformations. Subsequently, non-uniformity correction using T1w and T2w images was applied [109]. Preprocessed images were nonlinearly registered to MNI152 space and cortical surfaces were extracted using FreeSurfer 5.3.0-HCP [110–112], with minor modifications to incorporate both T1w and T2w [18]. Cortical surfaces in individual participants were aligned using MSMAll [113,114] to the hemisphere-matched conte69 template [115]. T1w images were divided by aligned T2w images to produce a single volumetric T1w/T2w image per subject (Glasser and Van Essen, 2011). Notably, this contrast nullifies inhomogeneities related to receiver coils and increases sensitivity to intracortical myelin.

For rs-fMRI, the timeseries were corrected for gradient nonlinearity and head motion. The R-L/L-R blipped scan pairs were used to correct for geometric distortions. Distortion-corrected images were warped to T1w space using a combination of rigid body and boundary-based registrations [116]. These transformations were concatenated with the transformation from native T1w to MNI152, to warp functional images to MNI152. Further processing removed the bias field (as calculated for the structural image), extracted the brain, and normalised whole-brain intensity. A high-pass filter (>2000s FWHM) corrected the timeseries for scanner drifts, and additional noise was removed using ICA-FIX [117]. Tissue-specific signal regression was not performed [118,119]. We finally transformed these rs-fMRI to native space and sampled time-series at each vertex of the MSMAll-registered [113,114] mid-thickness cortical surfaces.

#### In vivo microstructure profile covariance (MPC) analysis

We estimated MPC in the *in vivo* dataset (MPC_MRI_) in the same manner as MPC_HIST_ with the only adjustment that intensity profiles were not corrected for *y*-coordinates; instead, the contrast reversal for T1w and T2w data was used to correct for inhomogeneity as part of the HCP minimal processing pipeline. We generated equivolumetric surfaces between the outer and inner cortical surfaces (see Equation (1)), and systematically sampled T1w/T2w values along 64,984 linked vertices from the outer to the inner surface across the whole cortex. In turn, MPC_MRI_ can be denoted as an extension of Equation (2), in which MPC_MRI_*(i,j)* for a given pair of nodes *i* and *j* is defined by:

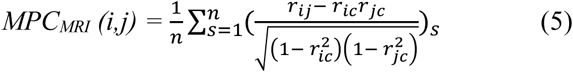

where *s* is a participant and *n* is the number of participants. We systematically evaluated matrix stability with 4 to 30 intracortical surfaces and selected 14 surfaces as the most stable solution (S10 Fig).

#### *In vivo* MPC_MRI_ *gradient: relation to the sensory-fugal gradient and the histological MPC gradient*

As for the histological data, diffusion map embedding derived a principal gradient (G1_MRI_) from the group average MPC_MRI_ matrix. Correspondence of the *in vivo* G1_MRI_ to the histological G1_HIST_ gradient was estimated via Spearman rank correlation between spatially matched nodes. To localise differences, we calculated the difference in rank of each node. To dispel potential confounds due to regional differences in the variance in curvature across participants, we calculated average curvature of each node for each subject, then estimated the correlation between node-wise standard deviation in curvature and the G1_HIST_ and G1_MRI_ difference map. As before, we assessed the contribution of level of laminar differentiation and cytoarchitectural class to the G1_MRI_ via multiple regression.

### Correspondence of microstructure and functional connectivity

Individual functional connectomes were generated by averaging preprocessed timeseries within nodes, correlating nodal timeseries and converting them to z scores. For each individual, the four available resting state scans were averaged at the matrix level. Then, a group average functional connectome was calculated across the *Discovery* cohort. Correlation coefficients were calculated between the group average functional connectome and the MPC_HIST_ and MPC_MRI_ matrices. The group average functional connectome was proportionally thresholded at 90% per row, transformed into a cosine similarity matrix, transformed into a normalised angle matrix, then diffusion map embedding was applied, producing G1_FUNC_. We calculated the correspondence of the G1_FUNC_ with G1_HIST_ and G1_MRI_ with Spearman rank correlations. The differences in the gradients were localised by comparing node ranks across the whole cortex and within functional communities. Differences were also calculated using vertex-wise gradients to ensure the effects were not confounded by averaging disparate microstructural or functional profiles within nodes. Again, we also assessed associations to across-subject variance in curvature and the G1_MRI_ and G1_FUNC_ difference map. Seven functional communities were mapped onto the conte69 surfaces from a previous parcellation [26] with nearest neighbour interpolation from fsaverage5, then nodes were assigned to the functional community most often represented by the underlying vertices. To aid interpretation of the modality-specific gradients, G1_MRI_ and G1_FUNC_ were discretised into fifty equally-sized bins and we calculated the proportion of each bin accounted for by each functional community, then performed seven paired t-tests contrasting average node rank of a functional community in G1_MRI_ and G1_FUNC_ across individuals.

We assessed which cognitive faculties are related to dissociations between G1_MRI_ and G1_FUNC_ using meta-analytic maps from Neurosynth [120]. Neurosynth (http://www.neurosynth.org) combines automated text mining and meta-analytical techniques to produce probabilistic mappings between cognitive terms and spatial brain patterns. We downloaded and surface projected meta-analytic z-statistic maps of 24 terms covering a wide range of cognitive functions, which correspond to the topic names defined by Margulies et al. (2016). We discretised G1_MRI_ and G1_FUNC_ into five-percentile bins and, for each term, calculated the mean z-statistic within each bin. From this, we deduced the centre of gravity of each term within gradient space and calculated the differences between the G1_MRI_ centre of gravity and G1_FUNC_ centre of gravity for each term using subject-specific gradients.

### Robustness of the MPC approach

#### *Individualised* MPC_MRI_ *gradients and relation to the individual-specific functional hierarchy*

Inter-individual consistency of G1_MRI_ was assessed in the *Replication* dataset. The MPC_MRI_ pipeline was deployed at an individual level, thus resolving individualised MPC_MRI_ matrices and gradients. Additionally, the diffusion map embedding was employed on functional connectomes to derive individual functional gradients [28]. To ensure the spatial correspondence of individual gradients, the individual gradients from the *Replication* dataset underwent Procrustes linear alignment to the *Discovery* dataset group average embedding. Individual cross-modal coupling was calculated as the Spearman rank correlation between G1_MRI_ and G1_FUNC_.

#### *Robustness of* G1_HIST_

We assessed the robustness of G1_HIST_ by altering pipeline parameters, repeating MPC_HIST_ generation and diffusion map embedding, then calculating the Pearson correlation of the modified G1_HIST_ with the original gradient. In particular, we evaluated variable matrix thresholds (*i.e.*, 70-95%, in steps of 1%), alternative surface number in which MPC_HIST_ matrix instability reached a local minima (*i.e.*, 54- and 91-surface solutions), the voxel resolution of the BigBrain volume (100-400 um), and spatial scale (*i.e.*, vertex-vs parcel-wise construction). For the latter, we correlated nodal gradient values with the median vertex within each parcel.

#### *Robustness of* G1_MRI_

We repeated the robustness procedures reported for the histological gradients in the *in vivo* dataset, including variation of thresholding level, parcellation usage and surface number. Here, the pipeline was repeated with 23 surfaces, pertaining to local minima in the *in vivo* MPC_MRI_ matrix instability.

#### *Independent replications of* G1_MRI_

We replicated the *in vivo* gradient in two independent datasets, based on two additional myelin sensitive magnetic resonance imaging contrasts.

##### a) Quantitative T1

We implemented the MPC approach on 17 healthy adults (5 females, mean±SD age=28.1±6.1, 2 left handed) for whom quantitative T1 relaxation time mapping (qT1) images were available. All participants gave informed consent and the study was approved by the local research ethics board of the Montreal Neurological Institute and Hospital. MRI data was acquired on a 3T Siemens Magnetom Prisma-Fit with a 64-channel head coil. A submillimetric T1-weighted image was acquired using a 3D-MPRAGE sequence (0.8mm isotropic voxels, 320×320 matrix, 24 sagittal slices, TR=2300ms, TE=3.14ms, TI=900ms, flip angle=9°, iPAT=2) and qT1 data was acquired using a 3D-MP2RAGE sequence (0.8mm isotropic voxels, 240 sagittal slices, TR=5000ms, TE=2.9ms, TI 1=940ms, T1 2=2830ms, flip angle 1=4°, flip angle 2=5°, iPAT=3, bandwidth = 270 Hz/px, echo spacing = 7.2ms, partial Fourier = 6/8). The combination of two inversion images in qT1 mapping minimises sensitivity to B1 inhomogeneities [121], and provides high intra-subject and inter-subject reliability [122].

Cortical surfaces were extracted from the T1-weighted scans using FreeSurfer 6.0 [110–112], and fourteen equivolumetric intracortical surfaces were generated [45]. qT1 was registered to Freesurfer native space using a boundary-based registration [116], and FreeSurfer native space was registered to standard conte69 space using Caret5 landmark-based registration [115]. We used the former to sample qT1 intensity values along the intracortical surfaces, and the latter to resample the evaluated surfaces to a common space with 64,984 matched vertices. As in the main approach, we averaged vertex-wise intensity profiles within 1012 nodes [38], computed pairwise partial correlations between nodal intensity profiles (controlling for the average intensity profile), kept only positive correlations, and log-transformed the result to produce a MPC_MRI-QT1_ matrix. Finally, we generated a group-average MPC_MRI-QT1_ matrix and applied diffusion map embedding. The similarity of G1_MRI-QT1_ to the original G1_MRI_ was measured with a node-wise Spearman rank correlation.

##### b) Magnetisation transfer

In an open dataset of 297 healthy young adults (149 female, mean±SD age=19.1±2.9; Kiddle et al., 2018), we studied intracortical depth profiles of magnetisation transfer (MT). MT is a validated measure of myelination [22], and was available in the form of average intensity values along eight equidistant intracortical surfaces (10% - 90% in 10% intervals) within 308 cortical regions (for further details see [92]). We similarly applied the MPC framework to the MT profiles, then averaged the MPC_MRI-MT_ matrix across the group and applied diffusion map embedding. We measured the Spearman rank correlation between G1_MRI-MT_ and the original G1_MRI,_ which was re-calculated with 308 matched cortical areas.

## Supplemental Information

**Figure S1.**
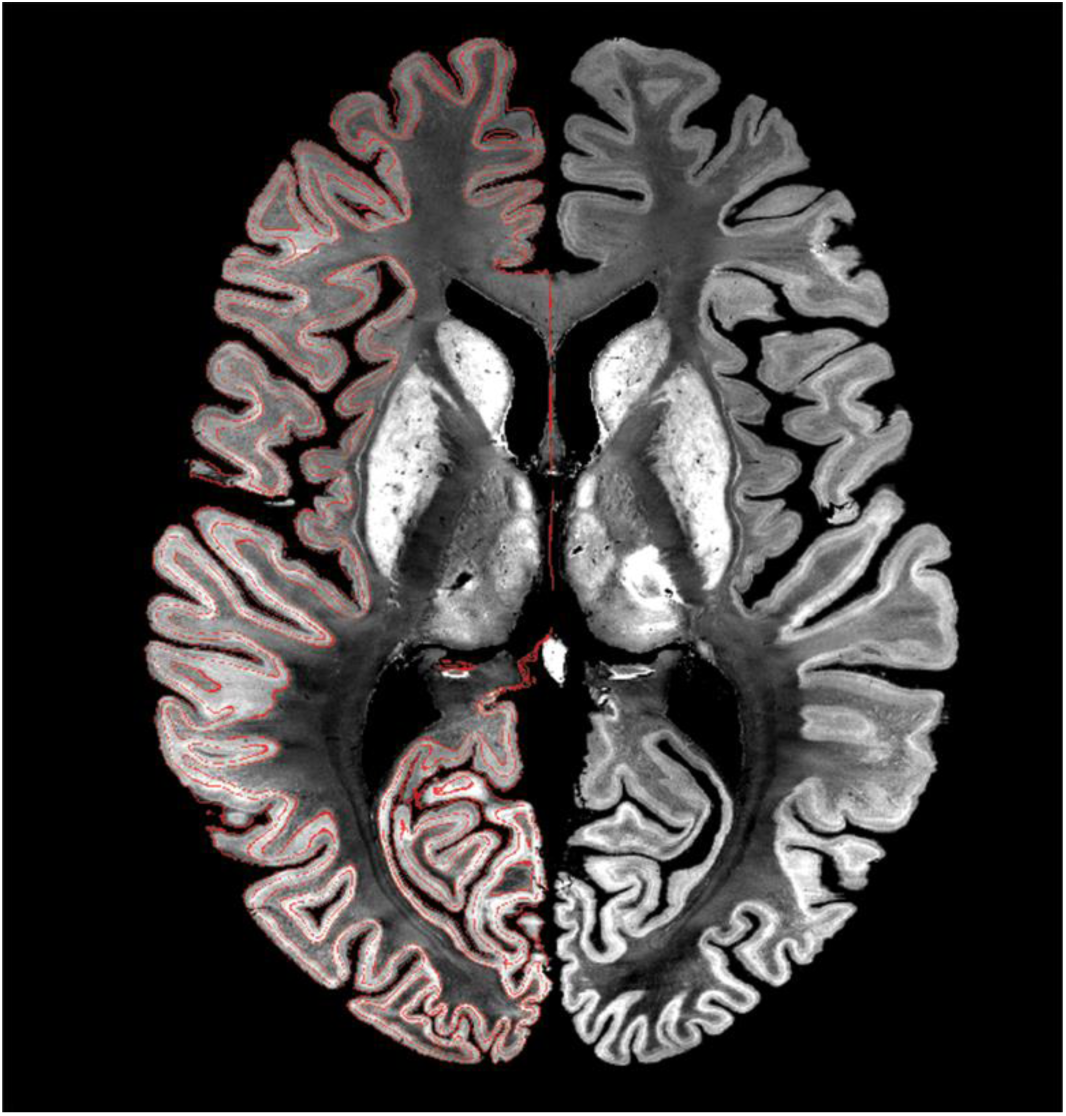
Horizontal view of the volumetric reconstruction of the “BigBrain” with pial, mid and white matter surfaces projected on the left hemisphere. Notably, we corrected for the linear relationship between intensity values and midsurface y-coordinate (r=-0.68, p<0.001), which existed due to coronal slicing and reconstruction of the BigBrain.

**Figure S2.**
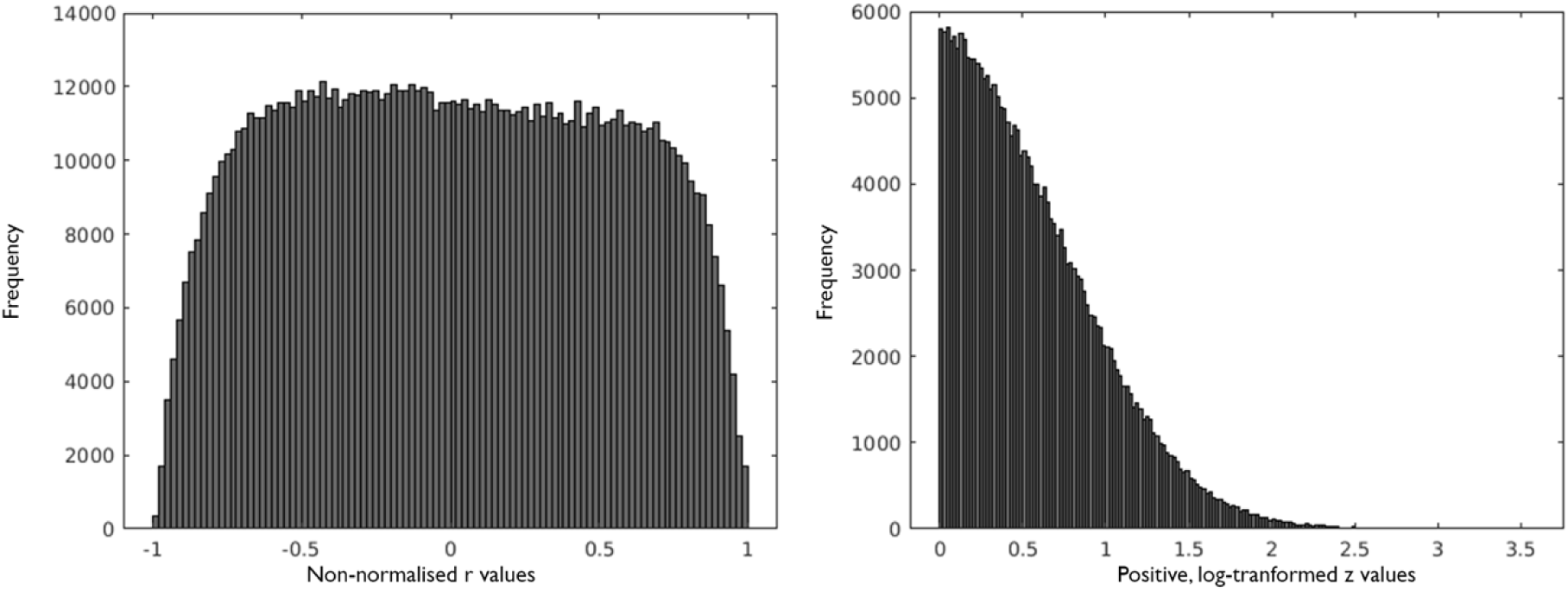
Distribution of values in MPC_HIST_ matrix. *(Left)* Frequency of r values calculated by Pearson product moment correlation coefficient of the nodal intensity profiles, controlling for the average intensity profile. *(Right)* Frequency of positive z values following log transformation of r values.

**Figure S3.**
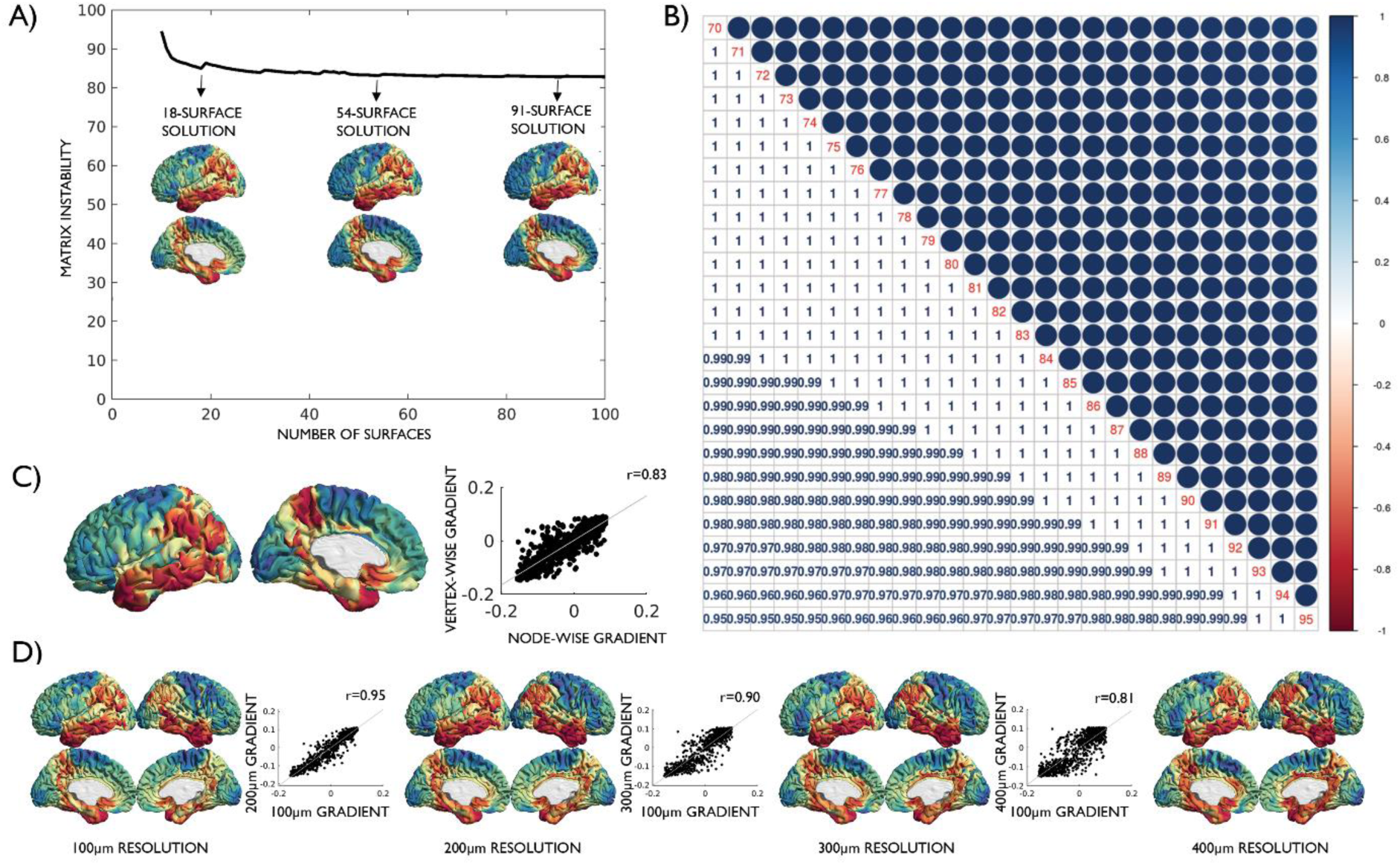
Robustness of G1_HIST_ to parameter variation. **(A)** MPC_HIST_ matrix instability using between 10 and 100 intracortical surfaces. G1_HIST_ was consistent regardless of the number of intracortical surfaces used, as shown by the strong spatial correlation of the 18-, 54- and 91-surface solutions (all r>0.97, all p<0.001). **(B)** Correlation matrix depicting the high correspondence of G1_HIST_ solutions with 70-95% row-wise matrix thresholding (0.95<r<1, all p<0.001). **(C)** Estimation of G1_HIST_ from 20488 vertices resulted in a consistent G1_HIST_ to the 1012 parcel construction pipeline (r=0.87, p<0.001). **(D)** The 200μm, 300μm and 400μm resolution BigBrain datasets were characterised as lower resolution replications, and G1_HIST_ was found to be highly correlated across these resolutions (all r>0.81, all p<0.001).

**Figure S4.**
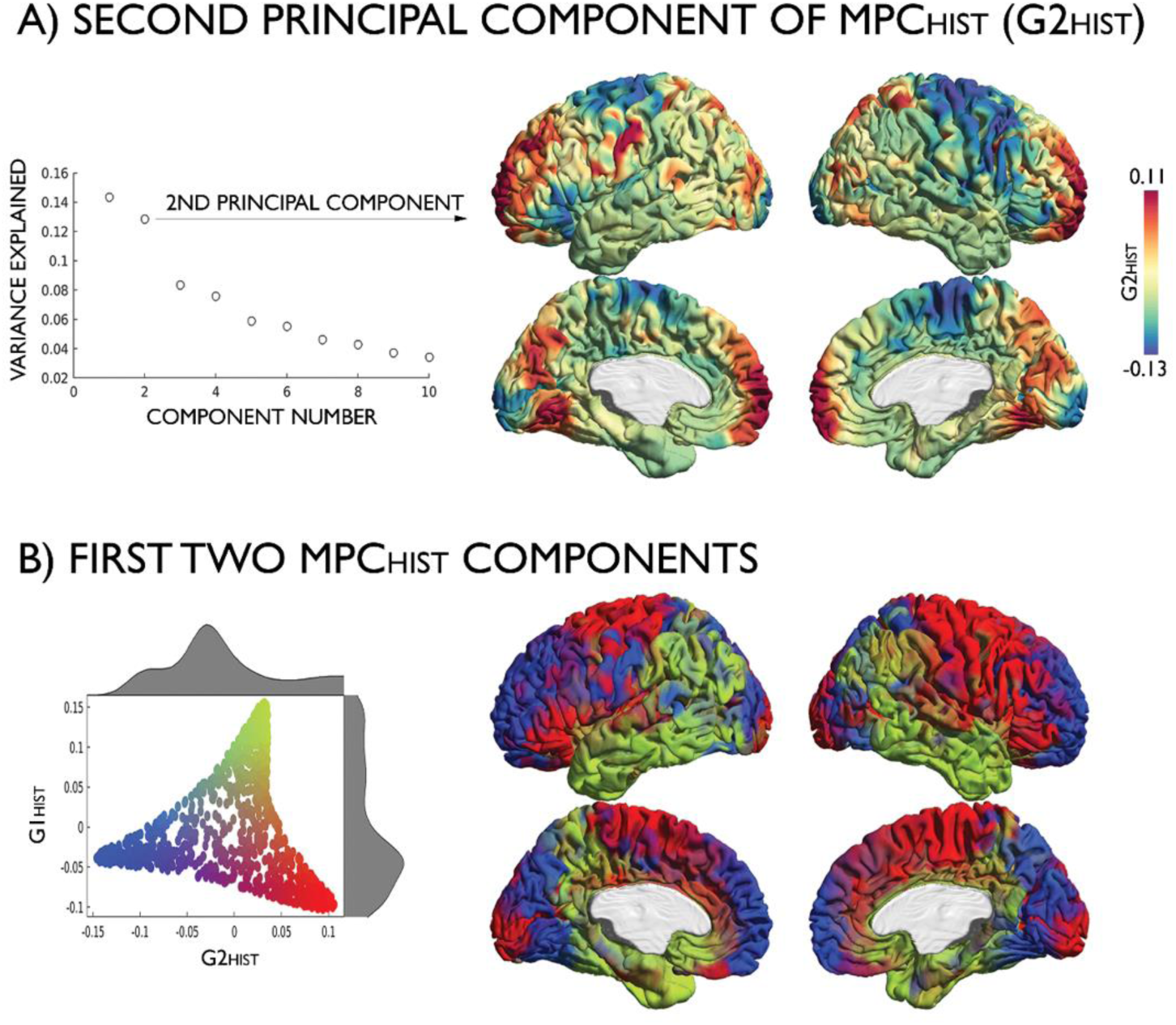
**(A)** The second principal component, accounting for 12.7% of variance in MPC_HIST_ components, is projected on the BigBrain midsurface. **(B)** Scatterplot depicting the first two embedding gradients, with corresponding probability density functions. The second gradient divides the lower-order areas of the first gradient, insomuch that somatomotor and primary visual areas *(red)* are separated from ventral prefrontal areas and secondary visual areas *(blue)*.

**Figure S5:**
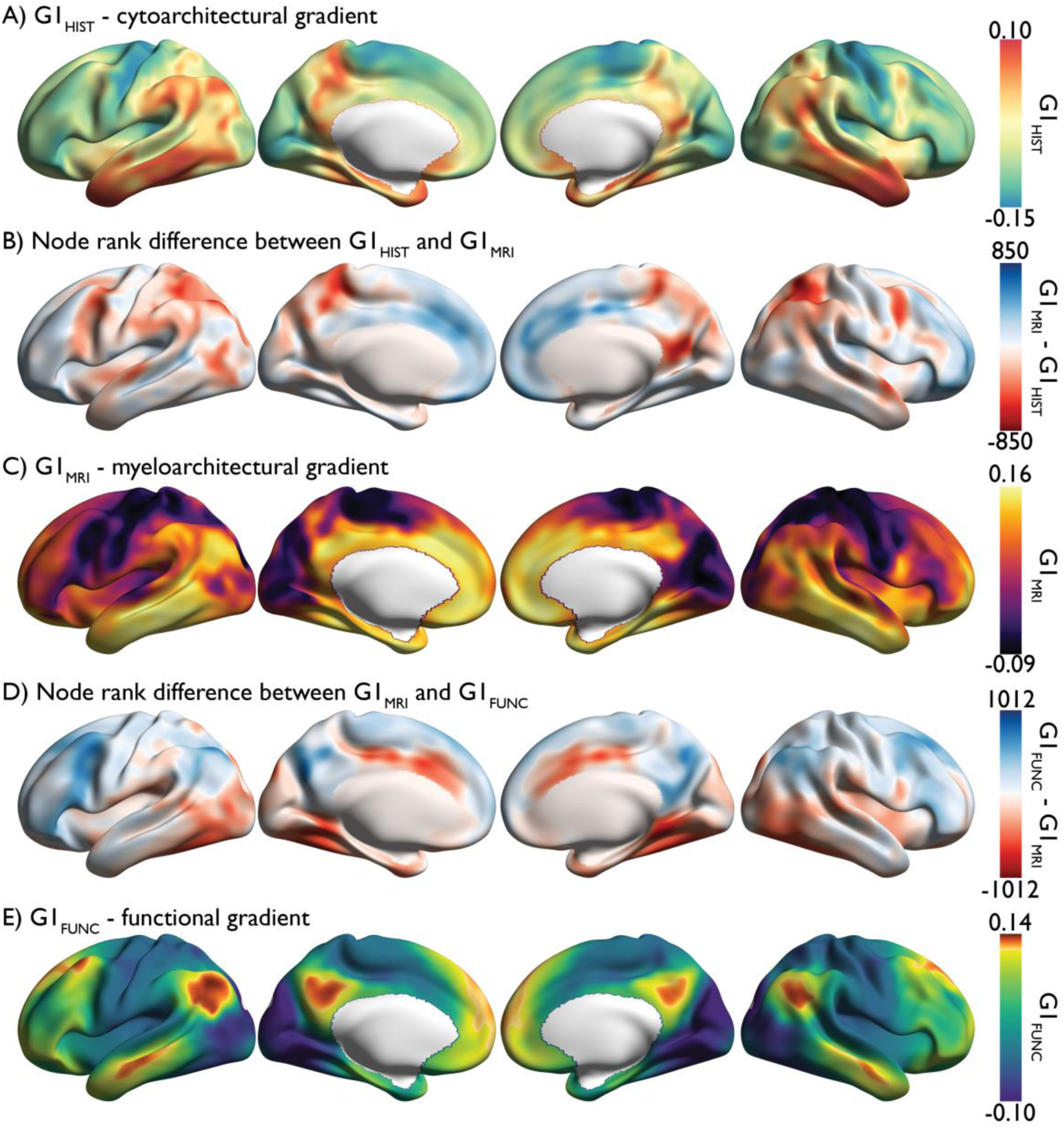
Principal gradients and node-rank differences projected on inflated cortical surfaces.

**Figure S6.**
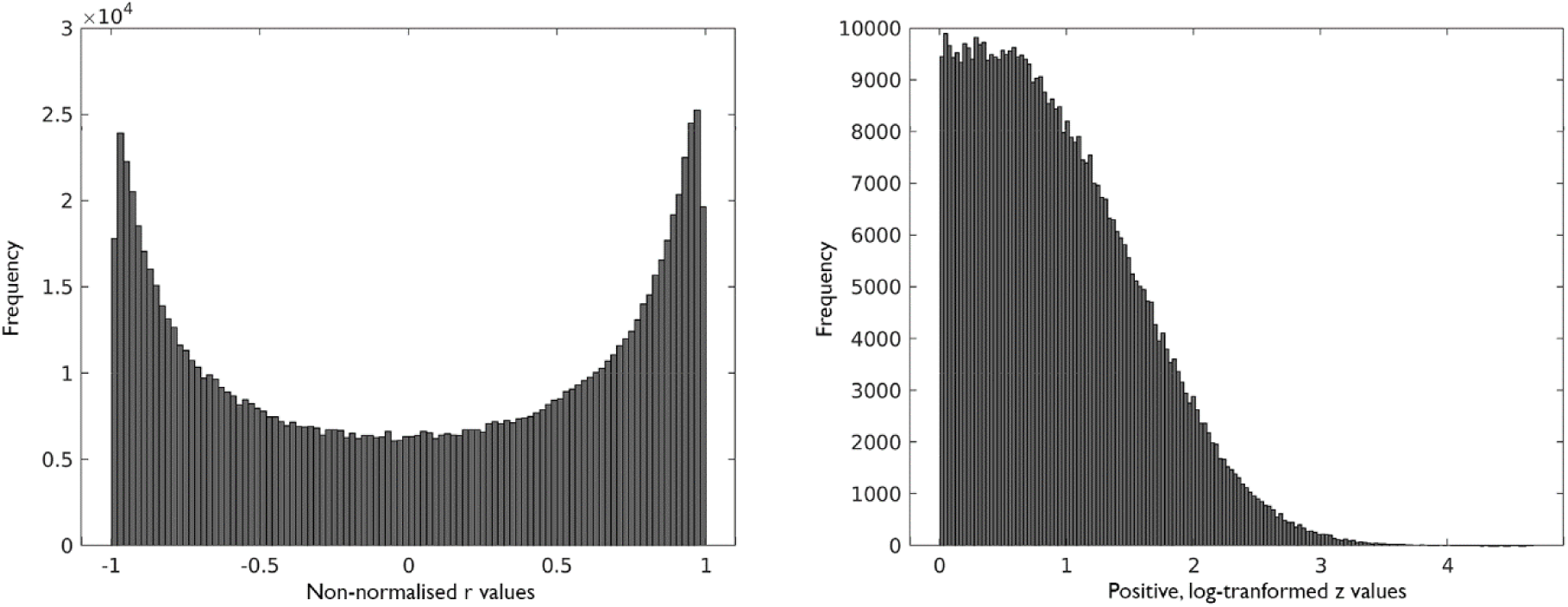
Distribution of values in MPC_MRI_ matrix. *(Left)* Frequency of r values calculated by Pearson product moment correlation coefficient of the nodal intensity profiles, controlling for the average intensity profile. *(Right)* Frequency of positive z values following log transformation of r values.

**Figure S7.**
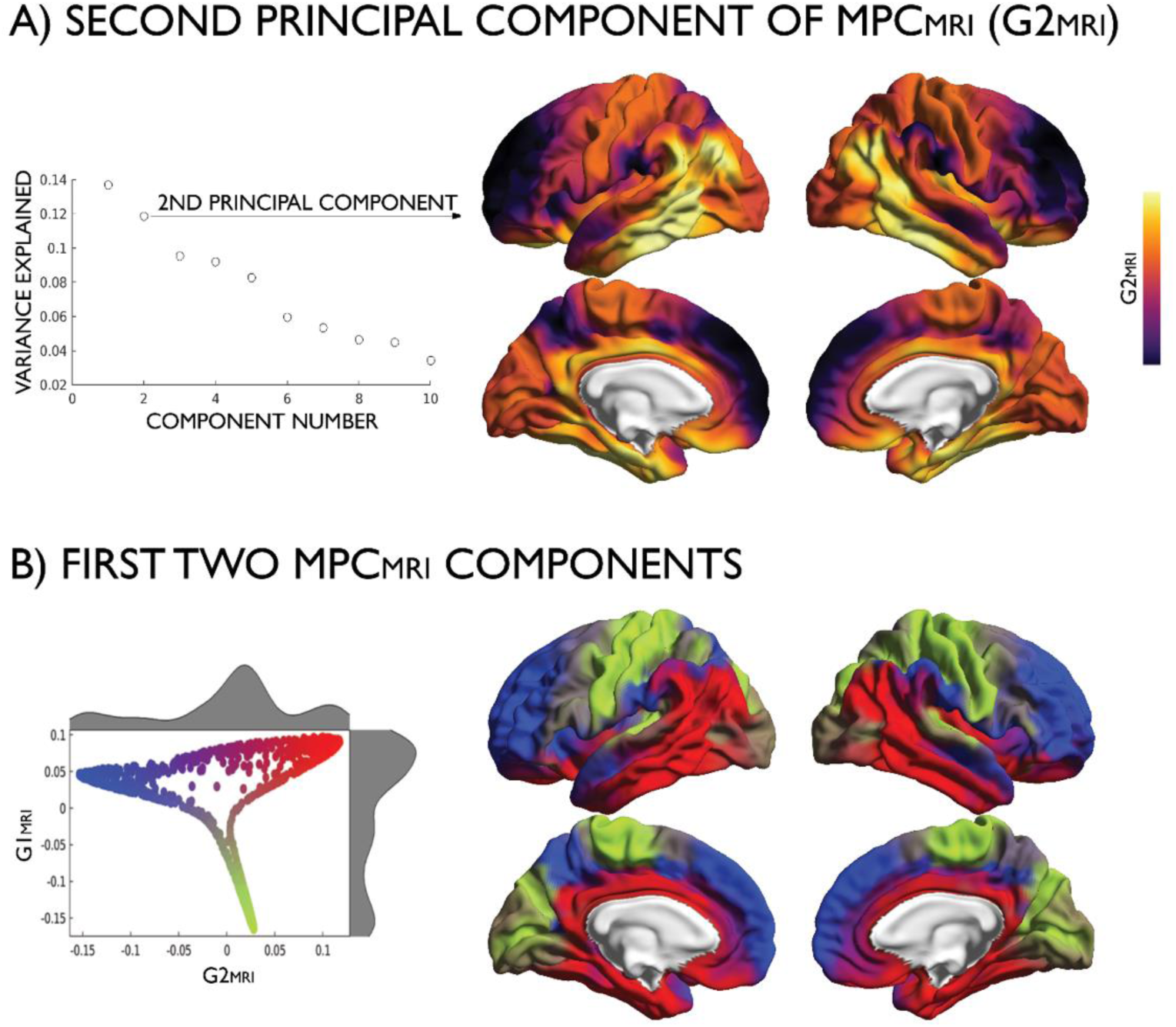
**(A)** The second principal component, accounting for 11.7% of variance in MPC_MRI_ components, projected on the conte69 midsurface. **(B)** Scatterplot depicting the first two embedding gradients, with corresponding probability density functions. The second gradient divides the higher-order areas of the first gradient, insomuch that the cingulate, orbitofrontal cortex and the inferior temporal gyrus *(red)* are separated from the prefrontal cortex, precuneus, temporo-parietal junction and superior temporal gyrus *(blue)*.

**Figure S8:**
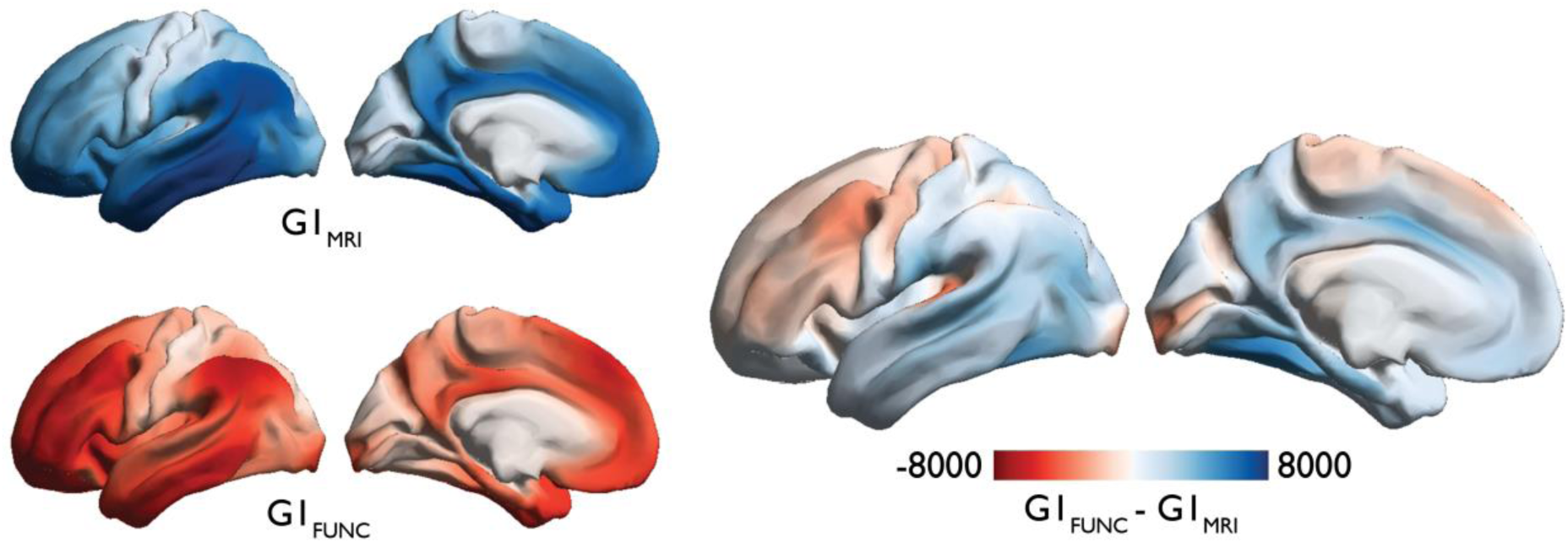
Differences in vertex ranks between G1_MRI_ *(blue)* and G1_FUNC_ *(red).*

**Figure S9.**
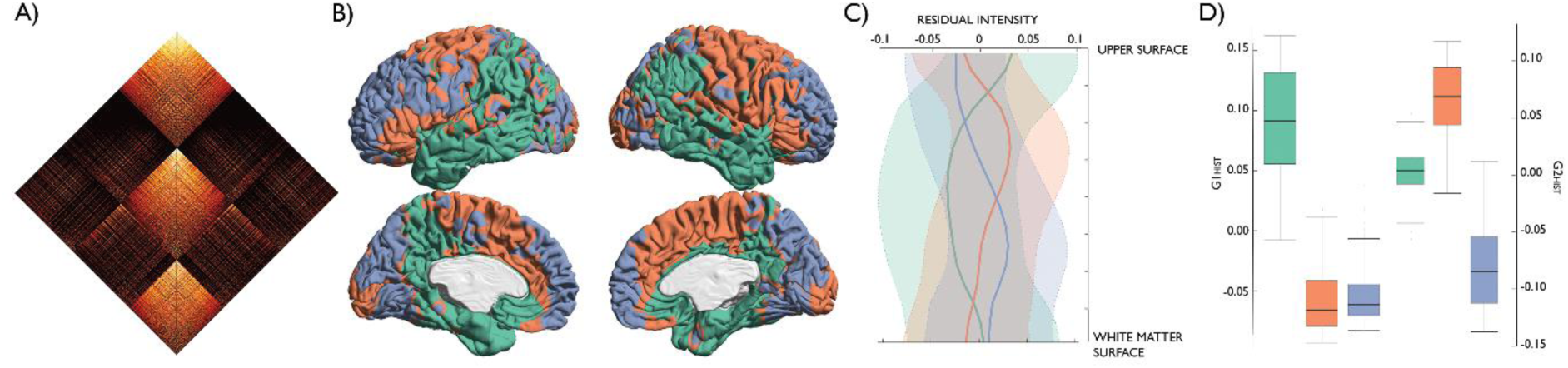
Community structure of the MPC_HIST_. **(A)** MPC_HIST_ matrix sorted by community membership. **(B)** Modular decomposition of MPC_HIST_ projected on the BigBrain midsurface. **(C)** Mean±sd of residual intensity profile for each module, after correction for the midsurface y-coordinate and demeaning. **(D)** Boxplot depicts the unique positions of modules along the first two principal gradients.

**Figure S10.**
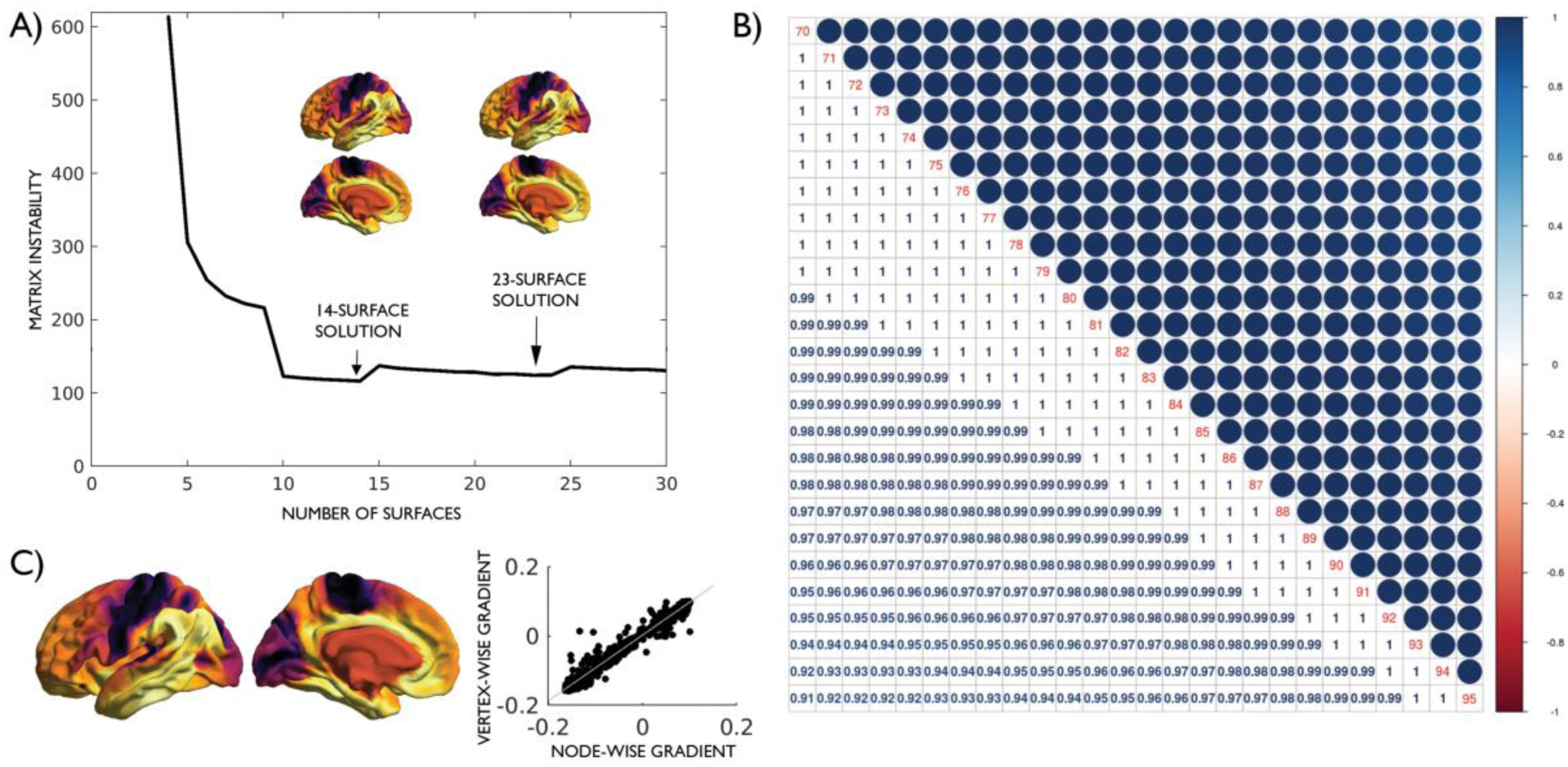
Robustness of G1_MRI_ to parameter variation. **(A)** MPC_MRI_ matrix instability using between 4 and 30 intracortical surfaces. G1_MRI_ was consistent regardless of the number of intracortical surfaces used, as shown by the strong spatial correlation of the 14- and 23-surface solutions (r=0.98, p<0.001). **(B)** Correlation matrix depicting the high correspondence of G1_MRI_ solutions with 70-95% row-wise matrix thresholding (0.91<r<1, all p<0.001). **(C)** Estimation of G1_MRI_ from 20464 vertices resulted in a consistent G1_MRI_ to the 1012 parcel construction pipeline (r=0.98, p<0.001).

**Figure S11.**
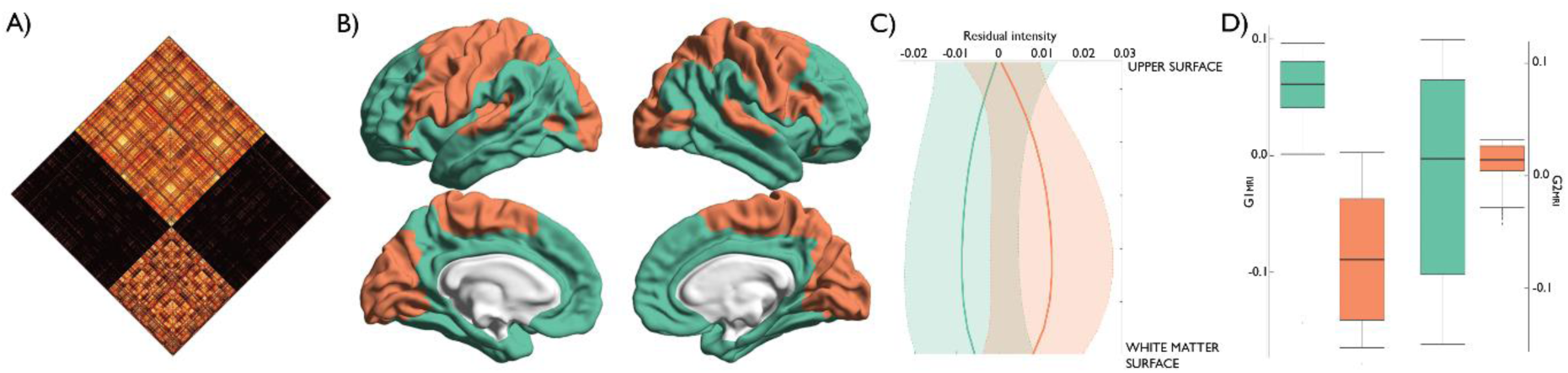
Community structure of the MPC_MRI_. **(A)** MPC_MRI_ matrix sorted by community membership. **(B)** Modular decomposition of MPC_MRI_ projected on the conte69 midsurface. **(C)** Mean±sd of residual intensity profile for each module, after demeaning. **(D)** Boxplot depicts the unique positions of modules on the first, but not the second, principal component.

**Figure S12.**
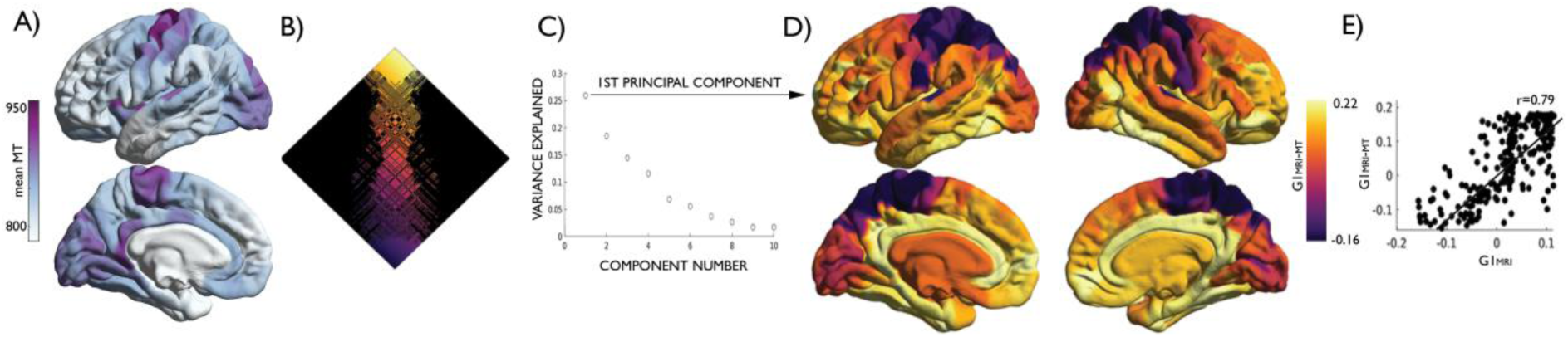
Independent replication of G1_MRI_, using magnetisation transfer (MT) data. **(A)** Mean MT intensity across subjects projected onto the cortical surface. **(B)** Gradient ordered normalised angle matrix. **(C)** Variance explained by embedding components. **(D)** First principal gradient (G1_MRI-MT_) projected onto conte69 midsurface. **(E)** Scatterplot depicting the strong correlation between the G1_MRI_ (reconstructed with the 308 parcellation scheme) and G1_MRI-MT_ (r=0.79, p<0.001).

**Figure S13.**
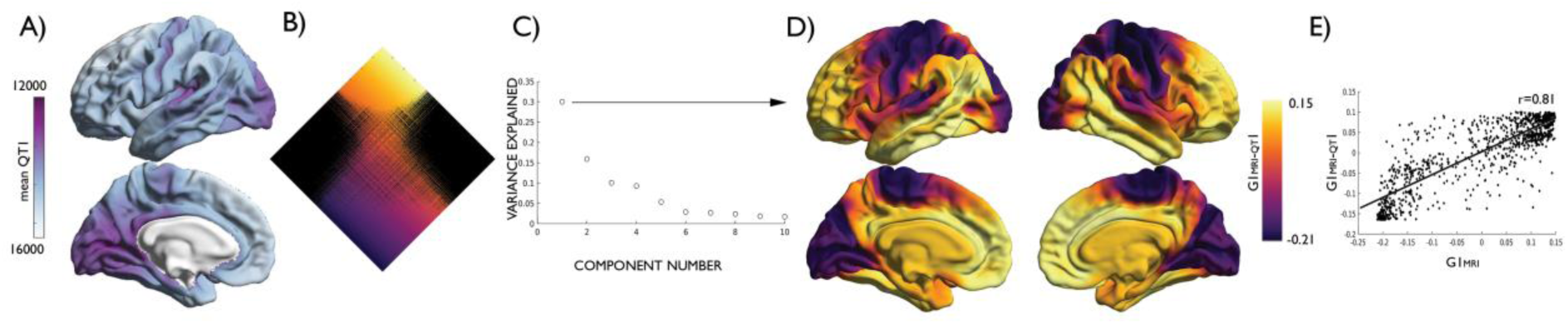
Independent replication of G1_MRI_, using quantitative T1 relaxometry (qT1) images. **(A)** Mean qT1 intensity across subjects projected onto the cortical surface. **(B)** Gradient ordered normalised angle matrix. **(C)** Variance explained by embedding components. **(D)** First principal gradient (G1_MRI-QT1_) projected onto conte69 midsurface. **(E)** Scatterplot depicting the strong correlation between the G1_MRI_ and G1_MRI-QT1_ (r=0.81, p<0.001).

**S1 Table:** Comparison of nomenclature and parcellation of cytoarchitectural groupings

| Mesulam, 2000 | Regional names <sup>1</sup> | Brodmann, 1909 <sup>1</sup> | Von Economo and Koskinas, 1925 |
| --- | --- | --- | --- |
| Idiotypic primary | Striate | 17 | OB, OC |
|  | Auditory | 41, 42 | TC |
|  | Somatosensory | 3a, 3b, 1, 2 | PA, PB, PC |
|  | Motor | 4,6 | FA |
| Modality-specific unimodal | Upstream peristriate | 18, 19 | OA |
|  | Inferotemporal | 20, 21, 37 | PH, TE |
|  | Superior temporal | 22 | TB, TD |
|  | Superior parietal lobule | 5, anterior 7 | Part of PE |
|  | Inferior parietal lobule | anterior 40 | Part of PF |
|  | Premotor | anterior 6, posterior 8, 44 | FB, FCBm |
| Higher-order heteromodal | Prefrontal cortex | 9, 10, 45, 46, 47, anterior 11, anterior 12, anterior 8 | FC, FD, FDdelta, FDT, FE |
|  | Posterior parietal | posterior 7, 39, 40 | PD, PG, parts of PE & PF |
|  | Lateral temporal | parts of 21 & 37 | Part TE |
|  | Parahippocampal | parts of 36 & 37 | TF |
| Paralimbic | Orbitofrontal cortex | posterior 11, posterior 12, 13 | FF, FG, FH, FJK, FLMH |
|  | Insula | 14, 15, 16 | IA, IB |
|  | Temporal pole | 38 | TG |
|  | Parahippocampal | 27, 28, 35 | HA, HB, HC |
|  | Cingulate | 23, 24, 25, 26, 29, 30, 31, 32, 33 | LA1, LA2, LC1, LC2, LC3, LD, LE |

<sup>1</sup> Ascribed by Mesulam, (2000)
*Note:* Mesulam (2013) defines transmodal cortex as areas without modality-specific input, thus including heteromodal and paralimbic isocortex.

**Table S2.** Individual predictors in a multiple regression model of G1_HIST_ by levels of laminar differentiation

|  | Estimate | Std. Error | t value | Pr(>\|t\|) |
| --- | --- | --- | --- | --- |
| Mesulam: idiotypic | 0.05651 | 0.004813 | 11.74 | 7.032e-30 |
| Mesulam: unimodal | -0.01105 | 0.003903 | -2.832 | 0.004723 |
| Mesulam: heteromodal | -0.005509 | 0.003352 | -1.644 | 0.1006 |
| Mesulam: paralimbic | -0.04945 | 0.006359 | -7.777 | 1.867e-14 |

**Table S3.** Individual predictors in a multiple regression model of G1_HIST_ by classes of cytoarchitecture

|  | Estimate | Std. Error | t value | Pr(>\|t\|) |
| --- | --- | --- | --- | --- |
| Von Economo: motor | 0.07335 | 0.008904 | 8.237 | 5.589e-16 |
| Von Economo: association<sup>1</sup> | -0.0005984 | 0.003267 | -0.1832 | 0.8547 |
| Von Economo: association<sup>2</sup> | -0.02703 | 0.00537 | -5.034 | 5.714e-07 |
| Von Economo: secondary sensory | 0.017 | 0.004712 | 3.607 | 0.000325 |
| Von Economo: primary sensory | 0.04783 | 0.008826 | 5.419 | 7.53e-08 |
| Von Economo: limbic | -0.06474 | 0.007497 | -8.636 | 2.32e-17 |
| Von Economo: insular | -0.02524 | 0.01259 | -2.004 | 0.04531 |

<sup>1</sup> Frontal and temporal association areas, displayed in yellow (**FIGURE 2**)<sup>1</sup> Parietal and superior temporal association, displayed in purple (**FIGURE 2**)

**Table S4.** Individual predictors in a multiple regression model of G1_MRI_ by levels of laminar differentiation

|  | Estimate | Std. Error | t value | Pr(>\|t\|) |
| --- | --- | --- | --- | --- |
| Mesulam: idiotypic | -0.06807 | 0.005734 | -11.87 | 1.76e-30 |
| Mesulam: unimodal | -0.03628 | 0.003094 | -11.73 | 7.847e-30 |
| Mesulam: heteromodal | 0.01026 | 0.003598 | 2.851 | 0.004444 |
| Mesulam: paralimbic | 0.09981 | 0.004438 | 22.49 | 8.103e-91 |

**Table S5.** Individual predictors in a multiple regression model of G1_MRI_ by classes of cytoarchitecture

|  | Estimate | Std. Error | t value | Pr(>\|t\|) |
| --- | --- | --- | --- | --- |
| Von Economo: motor | 0.1486 | 0.0086 | 17.28 | 1.251e-58 |
| Von Economo: association<sup>1</sup> | -0.01441 | 0.003227 | -4.467 | 8.831e-06 |
| Von Economo: association<sup>2</sup> | 0.001541 | 0.005227 | 0.2948 | 0.7682 |
| Von Economo: secondary sensory | -0.008014 | 0.004666 | -1.718 | 0.08619 |
| Von Economo: primary sensory | 0.0974 | 0.008251 | 11.8 | 3.57e-30 |
| Von Economo: limbic | -0.07451 | 0.007277 | -10.24 | 1.865e-23 |
| Von Economo: insular | -0.05544 | 0.01284 | -4.316 | 1.746e-05 |

<sup>1</sup> Frontal and temporal association areas, displayed in yellow (**FIGURE 2**)<sup>1</sup> Parietal and superior temporal association, displayed in purple (**FIGURE 2**)

**Table S6.** Statistical outcome of paired t-tests between G1_MRI_ and G1_FUNC_ within each functional community, taken across individuals.

| **Functional community** | **t-statistic** | **2.5% CI** | **97.5% CI** |
| --- | --- | --- | --- |
| Visual | -8.3215 | -76.998 | -47.508 |
| Somatomotor | -10.32 | -78.836 | -53.551 |
| Dorsal attention | -4.6335 | -35.345 | -14.249 |
| Salience | -29.099 | -202.85 | -177.11 |
| Limbic | -22.42 | -115.27 | -96.641 |
| Frontoparietal | 30.123 | 135.8 | 154.82 |
| Default mode | 54.445 | 165.76 | 178.21 |

